# Trial and error: a hierarchical modeling approach to test-retest assessment

**DOI:** 10.1101/2021.01.04.425305

**Authors:** Gang Chen, Daniel S. Pine, Melissa A. Brotman, Ashley R. Smith, Robert W. Cox, Simone P. Haller

## Abstract

The concept of *test-retest reliability* indexes the consistency of a measurement across time. High reliability is critical for any scientific study, but specifically for the study of individual differences. Evidence of poor reliability of commonly used behavioral and functional neuroimaging tasks is mounting. Reports on low reliability of task-based fMRI have called into question the adequacy of using even the most common, well-characterized cognitive tasks with robust population-level effects, to measure individual differences. Here, we lay out a hierarchical framework that estimates reliability as a correlation divorced from trial-level variability, and show that reliability estimates tend to be higher compared to the conventional framework that adopts condition-level modeling and ignores across-trial variability. We examine how estimates from the two frameworks diverge and assess how different factors (e.g., trial and subject sample sizes, relative magnitude of cross-trial variability) impact reliability estimates. We also show that, under specific circumstances, the two statistical frameworks converge. Results from the two approaches are approximately equivalent if (a) the trial sample size is sufficiently large, or (b) cross-trial variability is in the same order of magnitude as, or less than, cross-subject variability. As empirical data indicate that cross-trial variability is large in most tasks, this work highlights that a large number of trials (e.g., greater than 100) may be required to achieve precise reliability estimates. We reference the tools **TRR** and **3dLMEr** for the community to apply trial-level models to behavior and neuroimaging data and discuss how to make these new measurements most useful for current studies.

## 1 Introduction

The concept of test-retest reliability originated from the notion of inter-rater reliability, i.e., the measurement of agreement or consistency across different observers (Shrout and Fleiss, 1979). All statistics compress and extract information from data; test-retest reliability captures the degree of agreement or consistency across multiple measurements (rather than observers) of the same quantity (reaction time (RT), BOLD response, personality traits) under similar circumstances. Traditionally, reliability is assessed through the statistical metric of the intraclass correlation coefficient (ICC), which parses subject-associated variance and estimates the stability of that variance over time. In the current context we define test-retest reliability as a property of individual difference measures, and consider ICC the conventional statistical measure of reliability.

Assessment of reliability is critical for almost all data collected in scientific studies. Here we focus on one specific type of data structure common to behavioral and neuroimaging studies: subjects perform an experiment with different conditions aimed to probe specific processes; each condition is instantiated with many trials. For example, an experimenter might evaluate cognitive interference using the Stroop task (i.e., RT slowing due to conflicting information; MacLeod, 1991). The task includes two conditions, one where the name of the color (e.g., “blue”, “green”) and the color of the printed word match (i.e., congruent condition) and one where there is a mismatch between the print color and the color word (i.e., incongruent condition). The contrast between the two conditions renders an RT difference score (“contrast value”) per subject that indexes the ability to inhibit cognitive interference. If this task is completed twice by the same subjects, reliability across these two measures can be computed. Usually this measurement takes the form of ICC which represents the fraction of the total variance that can be attributed to inter-individual differences in the participants’ ability to ignore interfering information.

Assessment of reliability has always been part of rigorous questionnaire development; much less psychometric scrutiny has been applied to behavioral and imaging tasks until recently. Most concerningly, ICC estimates, especially those derived from contrast or subtraction scores (Meyer et al., 2017; Infantolino et al., 2018), now being reported appear unacceptably low for behavioral (around 0.5 or below; Hedge et al., 2018) and imaging tasks (less than 0.4; Elliott et al., 2020). This has cast doubt on the utility of these tasks in studies of individual differences. For neuroimaging tasks specifically, the extent to which common contrasts exhibit poor ICCs in brain regions of primary interest has been troubling. Task-based fMRI has been used for some time to examine individual differences in brain activation and inter-regional association as part of an important search for brain biomarkers of risk and disease. High reliability is a critical requirement for these usages.

Two aspects of the Stroop task example above are noteworthy and typical in modern test-retest reliability assessments of behavioral and imaging tasks: i) the experimenter seeks to assess the reliability of a contrast between two conditions, and ii) many trials are used as instantiations of each condition. While trials are clearly an important source of data variability, these are rarely included in the statistical model structure and certainly have not occupied a place in traditional reliability calculations via ICC. More broadly, modeling trial-level variance in general has not been widely practiced in neuroimaging (Chen et al.,2020). Most researchers estimate the trial-level effects and their contrasts through condition-level modeling (CLM). When condition-level effects are aggregated across trials, implicitly the researcher assumes no cross-trial variability or measurement error. If these assumptions were met, an ICC will adequately estimate the test-retest reliability. However, using a hierarchical framework with trial-level modeling (TLM), we show that, just as in behavioral studies (Rouder et al., 2019), trial-level variability is large, i.e., condition means are likely estimated with a significant amount of uncertainty for many common neuroimaging experiments. As per the law of large numbers in probability theory, the effect estimate for a condition should asymptotically approach the expected value with increased certainty as the number of trials grows. However, trial sample sizes would need to be very large to approach an estimate with high certainty, likely larger than in most current designs. As a result, the TLM-based framework, which explicitly accounts for trial-level variability, provides a more precise characterization of subject-associated variability and its stability over time. The limitations of CLM are not unique to the calculation of reliability but also apply to population-level analysis through CLM (Chen et al., 2020).

While the two modeling approaches, CLM and TLM, generate different reliability estimates, their results also converge under specific, rare circumstances. TLM-based reliability estimates are generally higher in the common scenario where the number of trials is low and/or cross-trial variability is high. We emphasize that the differences between the two modeling approaches is not conceptual. Rather, their similarities and differences originate from basic probability theory of sampling (e.g., the law of large numbers): when the trial sample size is sufficiently large or when the trial-to-trial variability is relatively small, CLM-based reliability estimates asymptotically approach their TLM-based counterparts.

### 1.1 Classical definition of ICC

In all comparisons with the ICC, we focus on the most common type, ICC(3,1), which quantifies the consistency (i.e., rather than absolute agreement) of an effect of interest between two repeated measures on a group of subjects (Shrout and Fleiss, 1979). Returning to our introductory example, suppose that the effect of interest is the contrast between incongruent and congruent conditions of the Stroop task. The investigator typically recruits *n* subjects who perform the Stroop task in two sessions while their RT is collected. Suppose that each of the two conditions is represented by *m* trials as exemplars in the experiment. The investigator usually follows the conventional two-level analytical pipeline. First, the original subject-level data *y_crst_* (*c* = 1,2; *r* = 1, 2; *s* = 1, 2, …,*n*; *t* = 1, 2,..,*m*) from the *s*-th subject during *r*-th repetition for the *t*-th trial under the *c*-th condition are averaged across trials to obtain the condition-level effect estimates 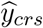. that are followed by contrasting the two conditions,

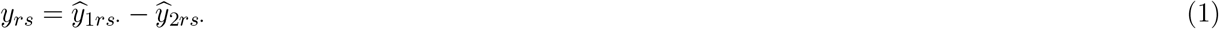

Then, at the population level, the condensed data *y_rs_* are fed into a CLM under a two-way mixed-effects ANOVA or LME framework with a Gaussian distribution,

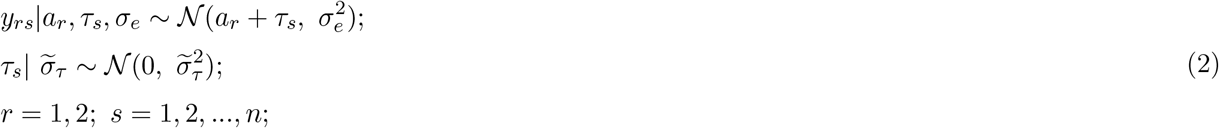

where *a_r_* represents the population-level effect during the *r*-th repetition, which is considered to be “fixed” under the conventional statistical framework; *τ_s_* is a “random” effect associated with *s*-th subject. The ICC under (2) is defined as

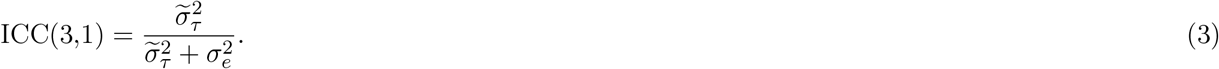

Such a two-level analytical pipeline, averaging across trials and contrasting between conditions followed by the ICC computation, is typically adopted for behavioral and neuroimaging data analysis. The residual variability *σ_e_* may appear to capture the within-subject cross-repetition variability; however, as we will detail later, “hidden” variability remains embedded in *σ_e_*.

The classical ICC can be viewed from two statistical perspectives. First, the total variance under the model (2) is partitioned into two components, *cross-subject variability* 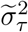 and *within-subject variability* 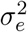. Thus, the ICC formulation (3) directly reveals the amount of cross-subject variability relative to the total variability. Alternatively, ICC can be viewed as the Pearson correlation of the subject-level effects between the two repetitions,

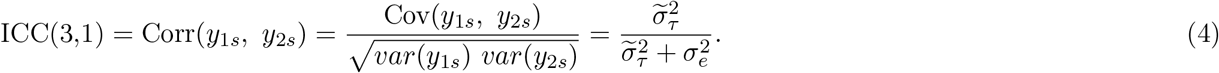

The ICC formulation is unique: unlike other generic *inter*class correlation where two variables are usually from two different classes (e.g., height and weight), ICC always involves two measures of the same class (e.g., the same type of effects *y*_1*s*_ and *y*_2*s*_). Further, homoscedasticity is implicitly assumed in the sense that the random variables *y*_1*s*_ and *y*_2*s*_ share the same variance across repetitions as shown in (4). This is distinct from a Pearson correlation, in which the two variables are usually heteroscedastic.

The computation of conventional ICCs is relatively straightforward. Using data from a publicly available dataset of the Stroop task (Study 1 in Hedge et al., 2018), we obtain a modest ICC(3,1) = 0.49 (Table 1) for the Stroop effect (i.e., contrast between the two conditions). The evidence for population-level effects is quite strong with a Stroop effect of *a*_1_ = 81 ms (95% uncertainty interval (72, 91)) and *a*_2_ = 59 ms (95% uncertainty interval (49, 69)) for the first and second repetition, respectively. As a comparison, the two conditions separately show a higher reliability with an ICC(3,1) of 0.72 and 0.69 for congruent and incongruent condition, respectively.

**Table 1:** Test-retest reliability (TRR) estimated through CLM and TLM for Study 1 data from the Stroop task (Hedge et al., 2018). RT (in ms) was recorded from 47 subjects, who completed two sessions of the Stroop task. With 240 trials per condition, RT ranged from 1 to 30830 ms; all data were used here without censoring. The variability ratio *R_v_* (defined in formulas (7) and (13)) and underestimation rate *R*_u_ (defined in formulas (6) and (12)) are indicators for the degree of underestimation through the CLM-based reliability estimation.

| Model | RT effect | Population Effects (ms) | | | Data Variability (ms) | | $R_v$<br>($R_u$) | TRR |
| --- | --- | --- | --- | --- | --- | --- | --- | --- |
|  |  | repetition | mean | 95% interval | subject | residual |  |  |
| model (2) | congruent | session 1 | 639 | (617, 660) | $\tilde{\sigma}_\tau$: 64 | $\sigma_e$: 39 | 4.1<br>(0.93) | 0.72 |
|  |  | session 2 | 607 | (585, 629) |  |  |  |  |
| CLM | incongruent | session 1 | 720 | (700, 744) | $\tilde{\sigma}_\tau$: 71 | $\sigma_e$: 47 | 3.0<br>(0.96) | 0.69 |
|  |  | session 2 | 666 | (641, 691) |  |  |  |  |
| ICC(3,1) | incongruent<br>vs congruent | session 1 | 81 | (72, 91) | $\tilde{\sigma}_\tau$: 23 | $\sigma_e$: 24 | 12.6<br>(0.61) | 0.49 |
|  |  | session 2 | 59 | (49, 69) |  |  |  |  |
| LME (5) | congruent | session 1 | 639 | (617, 660) | $\sigma_{\tau_1}$: 71 | $\sigma_0$: 300 | - | 0.78 |
| | | session 2 | 607 | (584, 629) | $\sigma_{\tau_2}$: 74 | | | |
| TLM | incongruent | session 1 | 720 | (693, 746) | $\sigma_{\tau_1}$: 89 | $\sigma_0$: 250 | - | 0.73 |
| | | session 2 | 666 | (643, 689) | $\sigma_{\tau_2}$: 77 | | | |
| LME (11) | incongruent<br>vs congruent | session 1 | 81 | (71, 92) | $\sigma_{\tau_1}$: 27 | $\sigma_0$: 276 | - | 1.0 |
| | | session 2 | 59 | (50, 68) | $\sigma_{\tau_2}$: 18 | | | |
| TLM | average of<br>2 conditions | session 1 | 679 | (656, 703) | $\sigma_{\tau_1}$: 79 | | - | 0.74 |
| | | session 2 | 636 | (614, 659) | $\sigma_{\tau_2}$: 75 | | | |
| BML (16) | incongruent<br>vs congruent | session 1 | 36 | (28, 42) | $\sigma_{\tau_1}$: 7 | $\sigma_0$: 75 | - | 0.71 |
| | | session 2 | 27 | (22, 33) | $\sigma_{\tau_2}$: 7 | | | |
| exGaussian | average of<br>2 conditions | session 1 | 670 | (656, 684) | $\sigma_{\tau_1}$: 34 | | - | 0.68 |
| | | session 2 | 643 | (631, 659) | $\sigma_{\tau_2}$: 28 | | | |

### 1.2 Separation between population effects and test-retest reliability

It is conceptually important to differentiate effects at different hierarchical levels (e.g., population, subject, and trial). Population-level effects are of general interest as researchers hope to generalize from specific subject samples to a hypothetical population. Population-level effects are captured through terms (usually called “fixed” effects) such as condition-level effects *a_r_* at the population level in the LME model (2). In contrast, lower-level (e.g., subject, trial) effects are mostly of no interest to the investigator since the samples (e.g., subjects and trials) are simply adopted as representatives of a hypothetical population pool, and are dummy-coded and expressed in terms such as subject-level effects *τ_s_* in the LME model (2).

It is important to recognize the dissociation between population effects and test-retest reliability. From the modeling perspective, the relationship between the population and subject level can loosely described as “crossed” or “orthogonal”: the subject-specific effects *τ_s_* in the LME model (5) are “perpendicular to” the population-level effects of overall average *a_r_* per repetition in the sense that the former are fluctuations relative to the latter. Even though cross-subject variability is the focus in the context of test-retest reliability, a popular misconception is that strong population-level effects are necessarily associated with high reliability as long as sample sizes are large. Fröhner et al. (2019) demonstrated the dissociation between the strength of population-level effects and the extent to which individual differences were reliable. Fig. 1 illustrates four extreme scenarios on a two-dimensional continuous space of population-level effects and reliability: one may have strong population-level effects with high (Fig. 1A) or low (Fig. 1B) reliability; contrarily, it is also possible to have weak population-level effects accompanied by high (Fig. 1C) or low (Fig. 1D) reliability.

**Figure 1:**
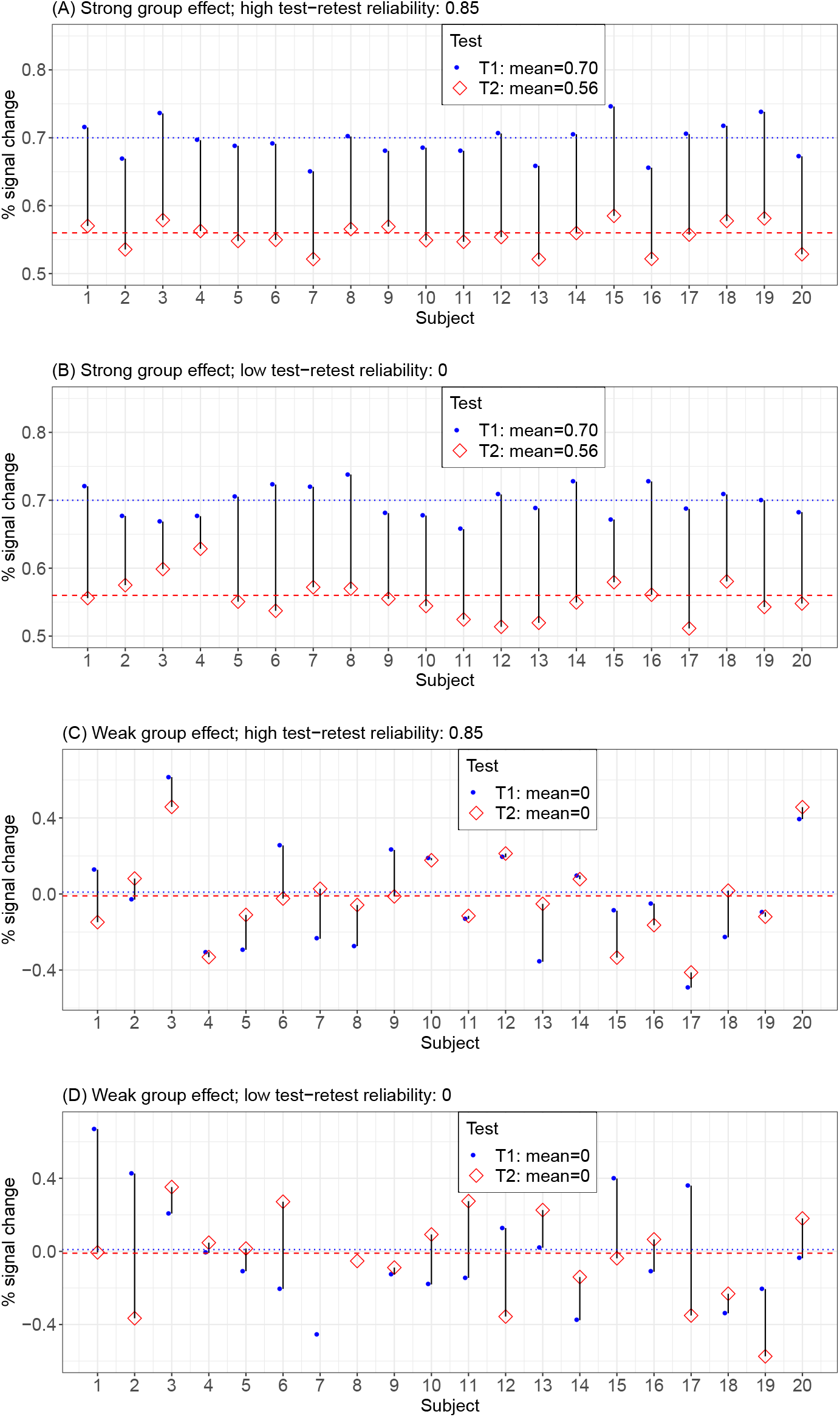
Separation between test-retest reliability and population-level effects. Four scenarios are illustrated to demonstrate that the reliability is not necessarily tied to the strength of population effects. Hypothetical data are based on a contrast between two conditions in a brain region: 20 subjects were scanned at two separate time points (T1: blue filled circle; T2: red empty diamond). Data were randomly drawn from a bivariate Gaussian distribution. The population effects are easy to observe via the colored horizontal lines. Reliability can be observed by assessing the proportion of subjects for which the two testing scores are on the same side (above or below) of their respective population average. With strong population effects, reliability can be high (A) or low (B); weak population effects may come alongside high (C) or low (D) reliability. 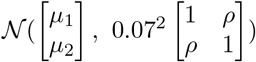 for each of the 20 subjects with (A) *μ*_1_ = 0.70, *μ*_2_ = 0.56, *ρ* = 0.85, (B) *μ*_1_ = 0.70, *μ*_2_ = 0.56, *ρ* = 0, (C) *μ*_1_ = *μ*_2_ =0, *ρ* = 0.85, and (D) *μ*_1_ = *μ*_2_ =0, *ρ* = 0.

## 2 Methods: assessing test-retest reliability through trial-level modeling

### 2.1 Motivation for adopting a new framework

The motivation for developing a hierarchical modeling approach is to derive more precise reliability estimates by modeling trial-level variability, thereby reflecting the data structure and generative mechanisms in our model. The conventional CLM-based approach assumes that all trials have the same effect; this assumption is implicit in the aggregation of trial-level measures through averaging for psychometric data or through a regressor with the same response across all trials (Fig. 2a-c). The condensation of all trials into a single estimate for the condition-level effect (without any uncertainty information) is used as input for reliability (and population-level) effect estimation. In contrast, a more precise approach is to capture the individual effects through TLM with one regressor per trial (Fig. 2d,e). We aim to address two questions, 1) What is the impact of modeling trial-level effects vis-a-vis CLM-based reliability estimation? 2) Under what circumstances do results from the two frameworks converge?

**Figure 2:**
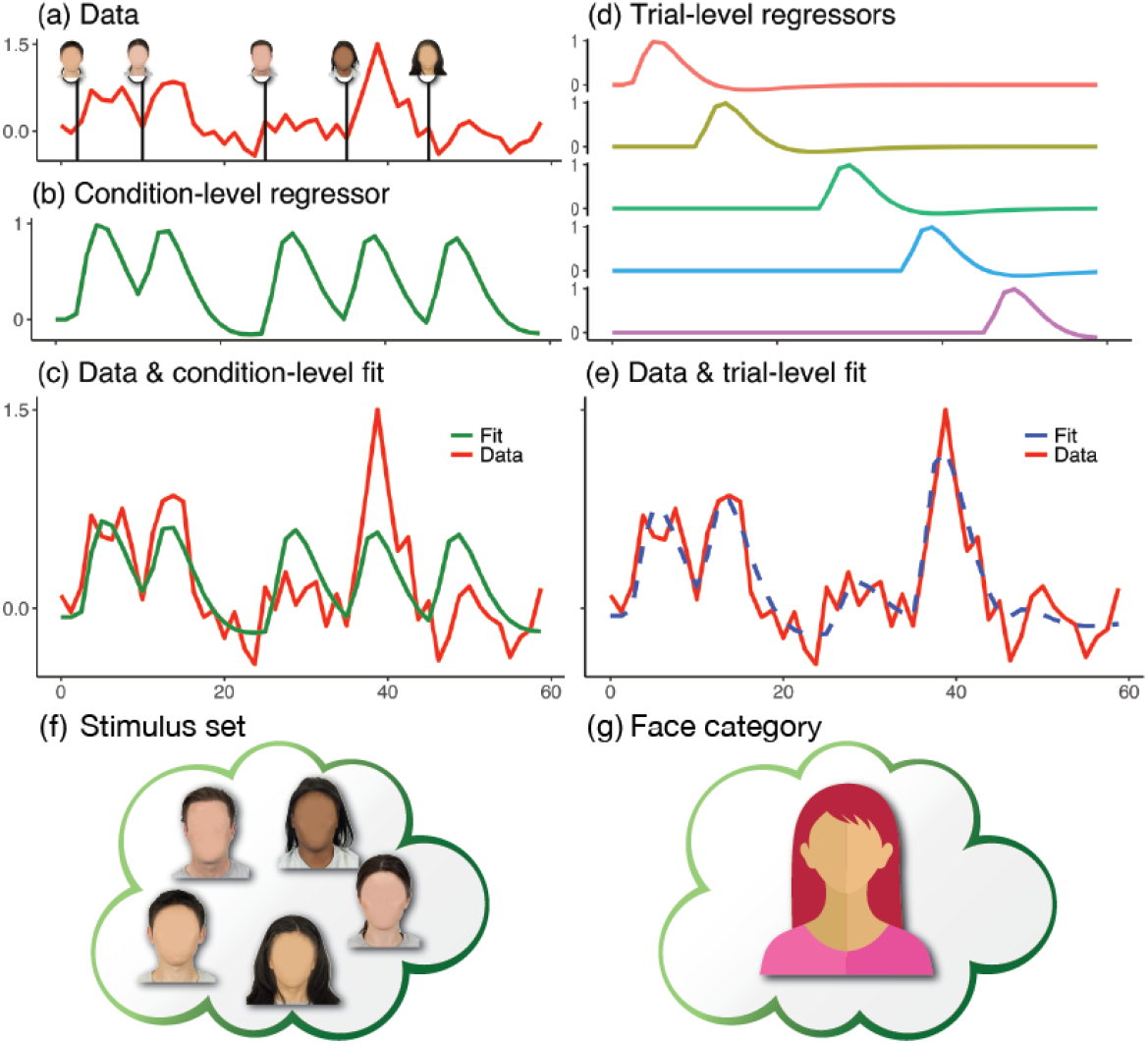
Condition-versus trial-level modeling. Consider an experiment with five face trials. (a) Hypothetical time series. (b) The conventional modeling approach assumes that all trials lead to the same response, so one regressor is employed. (c) Condition-level effect (e.g., in percent signal change) is estimated using the regressor fit (green) through condition-level modeling. (de) Trial-level modeling employs a separate regressor per trial, improving the fit (dashed blue). (f-g) Technically, the condition-level modeling does not allow for generalization to a face category. Note that modeling trial-level variance can be done both in terms of regular population-level models and, as described here, for reliability assessment.

To address these two questions, we are specifically interested in accounting for cross-trial variability in our hierarchical modeling. Trials are related to condition-level effects the same way participants are related to population-level effects. While trials are expected to provide robust estimation for condition-level effects, at the same time, variability associated with trials are largely of no interest; thus, in practice, the data are typically collapsed across trials and further flattened across conditions as illustrated in the data reduction formula (1) under the conventional CLM framework. As a result, cross-trial variability is not represented in the model and the associated uncertainty is neither accounted for nor propagated in the conventional formula (2) in the ICC computation.

We will also address the limitations of the CLM-based framework. First, it can be difficult to obtain a measure of reliability uncertainty. Approximations for the variance of reliability estimates are available (Shoukri et al., 2004) that could provide a confidence interval, but these require large subject sample sizes (because of the reliance on the asymptotic property). Even though bootstrapping could be adopted to find a quantile interval, the approach is rarely utilized in practice due to its computational cost, especially for large datasets in neuroimaging. Second, the LME formulation assumes a Gaussian distribution and, when this assumption is violated (e.g., skewed data, outliers), parameter estimation through the optimization of a nonlinear objective function can become unstable or singular. Third, it is not possible to integrate measurement error within the conventional platform. Under specific circumstances, the input data may contain sampling errors. For instance, BOLD response as the effect of interest in neuroimaging is not directly collected, but instead estimated from a time series regression model. Including measurement error can calibrate and improve the model fit. For example, instead of censoring at an artificial threshold, outliers can be more effectively handled through down-weighting based on their large uncertainty (Chen et al., 2020). This modeling strategy has been widely adopted to achieve higher efficiency and robustness in traditional meta-analysis (Viechtbauer, 2005) as well as in neuroimaging population-level analysis (Worsley et al., 2002; Woolrich et al., 2004; Chen et al., 2012). Although these approaches exist, the most common pipelines still use subject-level effect estimates for the population-level model without considering the associated uncertainty information; in addition, condition-level effects are estimated through one regressor per condition, ignoring cross-trial variability.

### 2.2 LME framework for a single condition

We start with simply accommodating trial-level effects in reliability estimation for a single condition in the LME formulation. There are five levels involved in the hierarchical structure of a reliability dataset (Fig. 3): population, subject, repetition, condition and trial. As opposed to the common practice of collapsing the two lower levels (condition and trial), we directly utilize the trial-level effect estimates *y_rst_* of the condition (Chen et al., 2020), where *r, s* and *t* index repetitions, subjects and trials (*r* = 1,2; *s* = 1,2,…,*n; t* = 1,2,..,*m*). Specifically, we expand the LME model (2) and accommodate the trial-level effects *y*_rst_ as below,

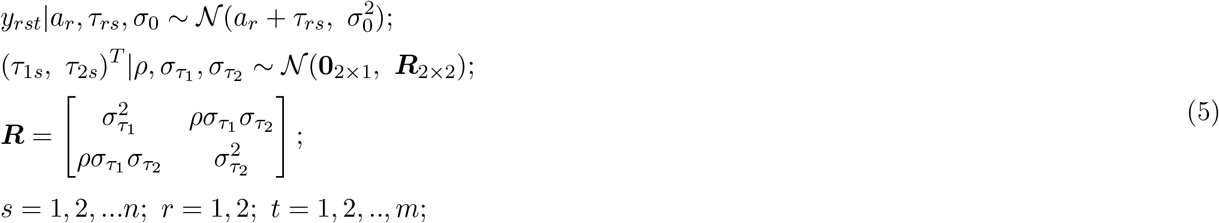

where *a_r_*, as in (2), represents the population-level effect during the *r*-th repetition, *τ_rs_* characterizes the subject-level effects during the *r*-th repetition, *σ*_0_ captures the cross-trial variability, and ***R*** is the variancecovariance matrix for the subject-level effects *τ_rs_* between the two repetitions. Usually *a_r_* are termed as the population-level (or “fixed”) intercepts while *τ_rs_* are the varying (or “random”) intercepts across subjects.

**Figure 3:**
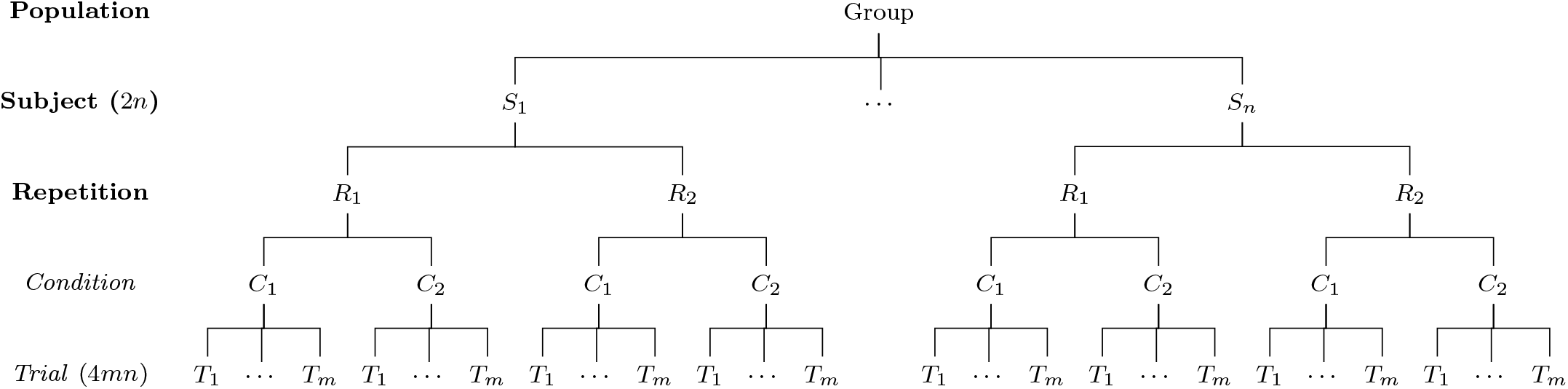
Hierarchical structure of test-retest data. Assume that, in a study with two repetitions, a group of n subjects are recruited to perform a task (e.g., Stroop) of two conditions (e.g., congruent and incongruent), and each condition is instantiated with m trials. The collected data are structured across a hierarchical layout of 5 levels (population, subject, repetition, condition and trial) with total *n* × 2 × 2 × *m* = 4*mn* data points at the trial level compared to 2n across-condition contrasts at the subject level.

The parameter *ρ* captures the correlation between the two repetitions for the subject-level effects *τ*_1*s*_ and *τ*_2*s*_. In fact, *ρ* represents the test-retest reliability of individual differences while the trial-level variability is explicitly characterized through *σ*_0_ in the model (5). The first line in the formulation (5) specifies the effect decomposition while the second line specifies the distributional assumptions of the terms. All the parameters including the reliability measure *ρ*, cross-subject variability *σ*_*τ*_1__ and *σ*_*τ*_2__, and cross-trial variability *σ*_0_ are numerically estimated through restricted maximum likelihood (Pinheiro and Bates, 2000). The inclusion of cross-trial variability *σ*_0_ into the LME model (5) precludes formulating reliability as a variance ratio as traditionally done in the formula (3). The reliability estimates for the Stroop task dataset based on the LME model (5) are shown in Table 1.

The relationship between ICC and the reliability metric *ρ* under the LME model (5) can be directly characterized. As is evident from Table 1, the CLM-based formulation tends to yield lower reliability estimates of a single condition compared to the TLM-based computations. For comparison, we temporarily assume homoscedasticity between the two repetitions: *σ*_*τ*_1__ = *σ*_*τ*_2__ = *σ*_*τ*_. The specific amount of underestimation can be revealingly expressed as a linear relationship (Appendix A),

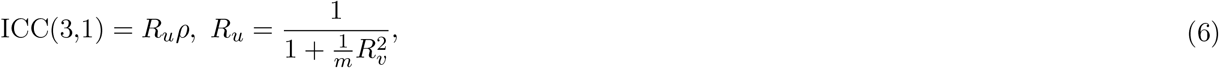

where the underestimation rate *R_u_* is dependent on the trial sample size *m* and the variability ratio *R_v_* (the magnitude of cross-trial variability relative to cross-subject variability)

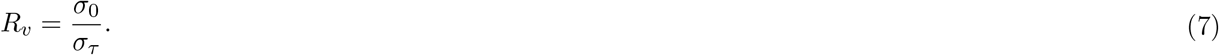

Both the differences and similarities between the two modeling frameworks can be clearly revealed through the ratio *R_u_*. First, the trial sample size m plays a crucial role; specifically, the degree to which the CLM-based estimates are lower than the TLM-based estimates decreases as the trial sample size increases. In contrast, the subject sample size *n,* on average, does not impact the difference in estimates. If the trial sample size m is so large that 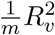 becomes negligible (i.e., *R_u_* ≈ 1), the two frameworks would render similar reliability estimates. Second, their divergence also depends on the variability ratio: the larger *R_v_*, the more severe the underestimation via the CLM. If cross-trial variability is roughly the same or smaller than its cross-subject counterpart (i.e., 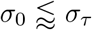) with a reasonable trial sample size *m* or if 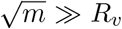, the difference between the two approaches is negligible (i.e., 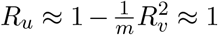) and CLM-based reliability estimates will asymptotically approach their TLM-based counterparts.

A correction could be applied to CLM-based reliability estimates post-hoc to adjust for trial-level variance if the variability ratio *R_v_* were known under the conventional framework (2). Specifically, the ICC formulation (3) could be adjusted to (Appendix A)

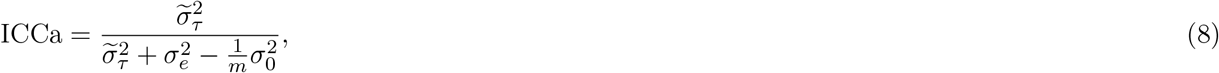

where 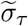 and *σ_e_* are the subject-level and residual variability under the conventional ICC formulation (2), while *σ*_0_ represents the cross-trial variability under the TLM-based LME formulation (5). As test-retest reliability is supposed to capture the correlation between the two condition-level effects (*τ*_1*s*_ and *τ*_2*s*_ in the model (5)) without the interference of trial-level fluctuations, it is desirable to explicitly account for the trial-level variability and to exclude it from the reliability formulation. In the same vein, the adjustment formula (7) accomplishes the same goal of removing the contamination of cross-trial variability through the term 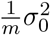. However, the adjusted ICC (8) is of little practical use because to estimate *σ*_0_, one would have to resort to the TLM-based LME formulation (5). In that circumstance, one might as well directly obtain the reliability through TLM.

That lower estimates are generated by the CLM-based formulation can be illustrated using the experimental data from the Stroop task (Hedge et al., 2018). The ICC(3,1) formulation generates values of 0.72 and 0.69 for the congruent and incongruent condition, respectively (Table 1). With the TLM-based LME model (5), we obtain a reliability estimate of 0.78 and 0.73 for each condition. In contrast, we could, per the formulation (7), obtain adjusted reliability estimates of 0.78 and 0.72 for the two conditions; the results are largely consistent with those from the TLM-based LME. The small amount of underestimation with *R_u_* = 0.93 or 0.96 was due to a large trial sample size (*m* = 240) and moderate variability ratios of R_v_ = 4.1 and 3.0.

### 2.3 Model comparisons through simulations

We adopt simulations to validate our theoretical reasoning in the previous subsection and to compare the two modeling frameworks in terms of a single condition. The numerical schemes for the simulations are presented in Appendix B with results shown in Fig. 4. The population-level effects *a_r_* under both CLM and TLM were robustly recovered (not shown). Findings and implications can be summarized as follows:

1. **The CLM-based approach leads to systematically lower reliability estimates compared to the TLM-based approach.** Simulations also confirmed the dependency of this difference in estimates on trial sample size *m* (Fig. 4B) and variability ratio *R_v_*. When cross-trial variability is much larger than cross-subject variability (*R_v_* » 1) the difference between the two is largest. When 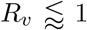 (top row, Fig. 4A), the simulated reliability values were successfully recovered for both formulations. When *R_v_* > 1 (second to fourth row, Fig. 4A), the bigger the ratio *R_v_*, the larger the discrepancy between the estimates from the two frameworks; as predicted, a linear relationship at the attenuation rate *R_u_* in (6) (green lines, Fig. 4A,B,C) emerged between ICC (red triangles) and the underlying reliability.
2. **The CLM-based estimates could be adjusted.** Once the cross-trial variability component 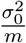 is explicitly accounted for in the ICC formulation as shown in (8), the adjusted ICCa generates estimates close to those derived from the TLM; the adjustment is quite successful when the sample size for both subjects and trials is 40 or above (second to fourth row, Fig. 4A).
3. **The subject sample size, on average, has no impact on the divergence in estimates between the two frameworks** (see also Shoukri et al., 2004; Noble et al., 2021). As shown in Fig. 4C, the degree of reliability attenuation by ICC (green line) relative to the TLM-based estimates (blue line) does not depend on the number of subjects n. However, as the subject number n increases, the precision of both reliability estimates improve.
4. **Achieving high precision of reliability estimates can be difficult.** The uncertainty associated with reliability estimates is monotonically associated with the reliability magnitude, variability ratio *R_v_*, and the sample sizes m and n for subjects and trials. Specifically, uncertainty reduces as (a) m or n increases, (b) reliability increases, or (c) *R_v_* becomes smaller. While both sample sizes influence reliability precision, with increases in m or n (from first to fourth column, Fig. 4) resulting in narrower error bars, the trial sample size m plays a more substantial role than the subject sample size n. In other words, large sample sizes for both subjects and trials (e.g., 100 or more when *R_v_* ≥ 10) may be required to achieve a reliability estimate with a reasonable precision, but the impact of the trial sample size m is slightly more impactful than the subject sample size n (Fig. 4B vs. Fig. 4C).
5. **Reliability is not associated with population-level effects.** The two sets of population-level parameters, (*a*_1_, *a*_2_) = (0, 0) and (1.0, 0.9), describe a scenario with no population effects and one with strong population effects. For both scenarios, simulation parameters were successfully recovered from the TLM-based LME formulation (5), and the two scenarios rendered very similar reliability patterns (note, only (*a*_1_, *a*_2_) = (0, 0) is shown in Fig. 4).
6. **LME modeling is susceptible to numerical instability.** The performance of the TLM-based LME formulation is acceptable, but suffers from numerical failures with a slight overestimation of reliability when cross-trial variability is much larger than cross-subject variability (*R_v_* » 1) (second to fourth row, Fig. 4A). In fact, the larger the ratio *R_v_*, the more severe the overestimation. Overestimation via the LME model occurred when the numerical optimizer was trapped at the boundary of *ρ* =1 with the convergence failure leading to a reliability estimate of 1.0. When *R_v_* > 10, numerical instability worsens substantially, consistent with the result from the Stroop task data in Table 1.

**Figure 4:**
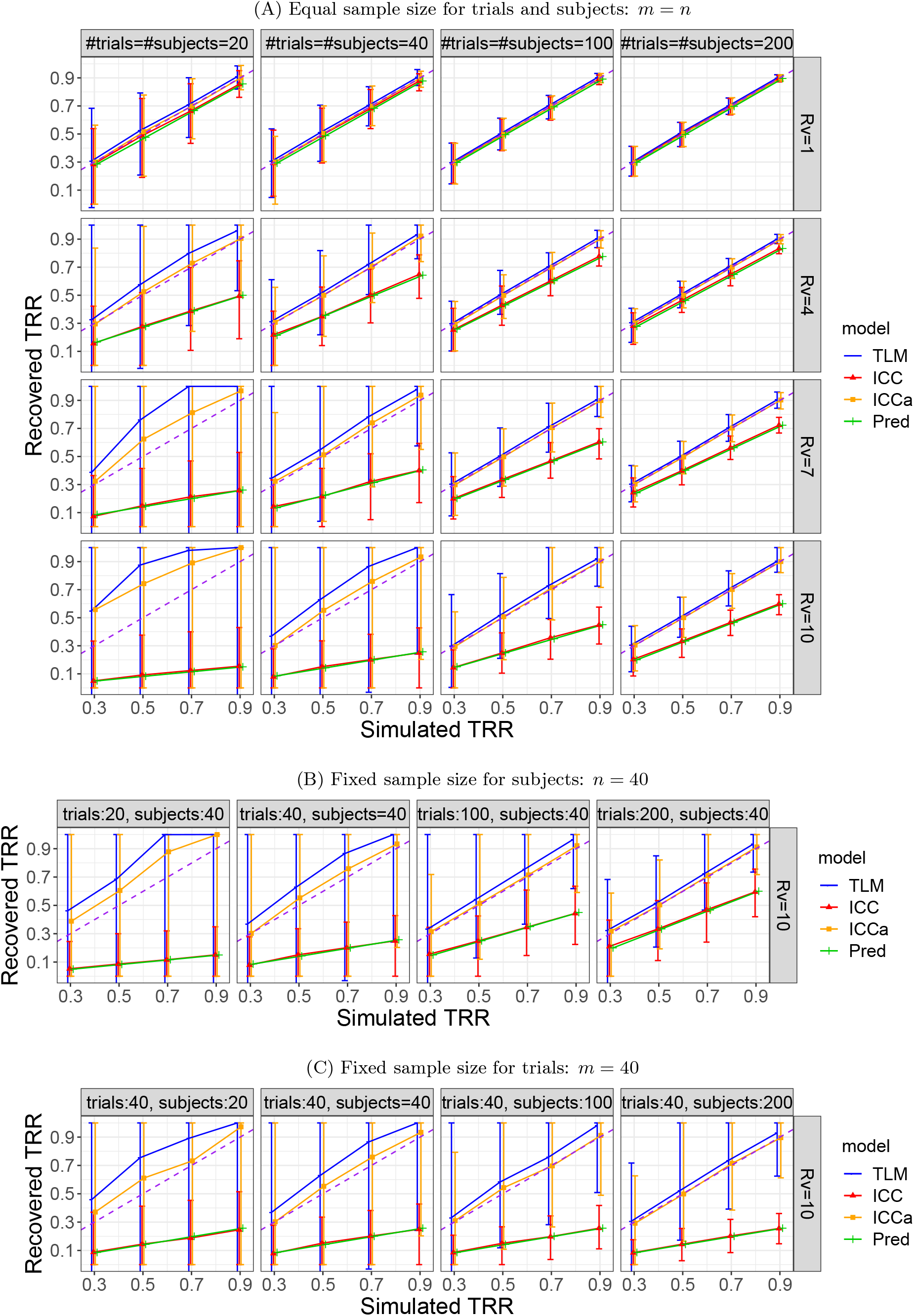
Simulation results for a single condition. The four columns correspond to subject and trial sample sizes, while the four rows code the variability ratio 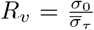. The *x*-and *y*-axis are the simulated and recovered reliability, respectively. Each data point is the median among the 1000 iterative simulations with the error bar showing the 90% highest density interval. The dashed purple diagonal line indicates an ideal retrieval of the parameter value based on the data-generating model (5). The green line shows the theoretically predicted ICC per the formula (6).

### 2.4 LME framework for a contrast between two conditions

Now we extend the LME formulation to the more common scenario of a contrast between two conditions. Suppose that we obtain the trial-level effects *y_crst_* of two conditions, where the four indices *c, p, s* and *t* code conditions, subjects, repetitions and trials. The LME formulation would read:

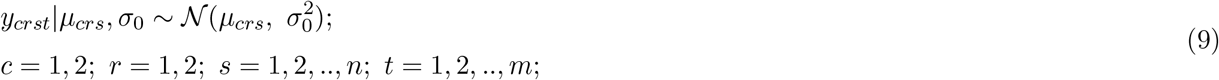

where *μ_crs_* is the *s*-th subject’s effect under the *c*-th condition during the *r*-th repetition, and *σ*_0_ captures the within-subject, within-repetition cross-trial variability. When the contrast is of interest, we need to parameterize the subject-level effects *μ_crs_* through dummy coding. Different methods exist for factor parameterization; we opt to quantify the two conditions through the following indicator,

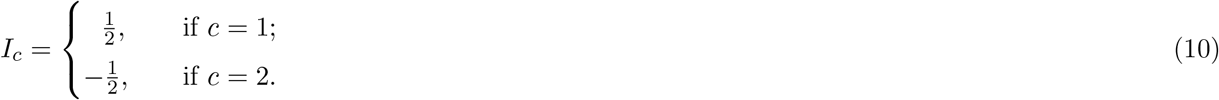

The subject-level effects *μ_crs_* are further integrated into the LME formulation (9) via,

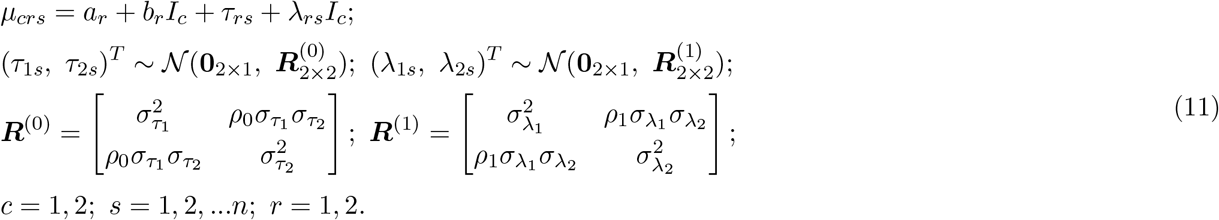

The variance-covariance matrices ***R***^(0)^ and ***R***^(1)^ characterize the relatedness across subjects between the two repetitions for the two parameter sets of (*τ*_1*s*_, *τ*_2*s*_)^*T*^ and (*λ*_1*s*_, *λ*_2*s*_)^*T*^, respectively. The assumption of independence between two parameter sets is discussed in Appendix C.

The correlation *ρ*_1_ embedded in the matrix ***R***^(1)^ captures the reliability for the contrast between the two conditions. With the two conditions coded through the indicator variable *I_c_* in (10), the contrast effects correspond to the slope terms (i.e., *b_r_* and *λ_rs_*) under the LME framework (11). Specifically, *b_r_* is the population-level contrast while *λ_rs_* codes the subject-specific contrast effects. In parallel to the single condition in (5) with the intercept interpretation, here *a_r_* and *τ_rs_* are also intercepts: the former is the population-level average between the two conditions during the *r*-th repetition while the latter indicates the condition average for the *s*-th subject during the *r*-th repetition.^1^ Just as the matrix ***R*** in (5) for a single condition, the matrix ***R***^(1)^ characterizes the relationships of subject-level contrasts *λ_rs_* between the two repetitions; thus, reliability can be similarly retrieved from the LME model (11) as the correlation *ρ*_1_ between the two varying slopes *λ*_1*s*_ and *λ*_2*s*_. As a byproduct, the correlation *ρ*_0_ embedded in the matrix ***R***^(0)^ for cross-subject varying intercepts *τ_rs_* captures the reliability for the average between the two conditions. For neueroimaging data analysis, whole-brain voxel-wise reliability for a contrast under the TLM-based LME framework (11) can be performed using the program **3dLMEr** (Chen et al., 2013) in AFNI.

What does explicitly modeling cross-trial variability under TLM reveal regarding the performance of the ICC formulation for a condition-level contrast? With the homoscedasticity assumption *σ_λ1_ = σ\_2_ = σ_λ_,* the extent of attenuation via the conventional ICC for a contrast is updated from the single effect case (6) to (Appendix D)

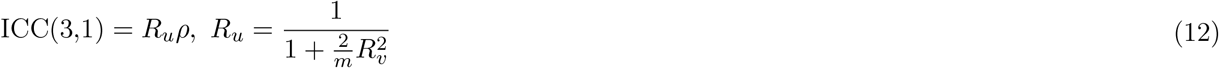

with the variability ratio *R_v_* similarly defined as before,

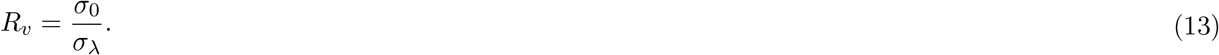

Likewise, one could adjust the conventional ICC by removing the impact of the cross-trial variance (Appendix D),

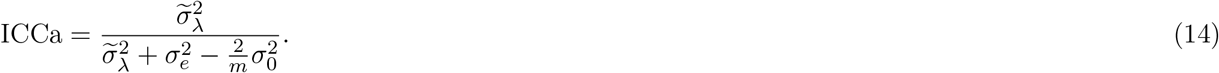

Compared to (6) and (8) for the case of a single condition, the number 2 occurs in the denominator of *R_u_* in (12) because there is twice the amount of data involved in the cross-trial variability for a contrast.

Table 1 compares reliability estimates for the condition-level contrast of the experimental Stroop data derived from the two frameworks. The ICC(3,1) value for the contrast was estimated as 0.49. For the corresponding TLM-based LME model, a reliability estimate of 1.0 was retrieved due to a singularity problem with the parameter *ρ*_1_ numerically degenerating at the boundary. With *σ*_0_ = 0.276, we adjusted the reliability to be 1.18 per (14). This uninterpretable value originated because the adjusted cross-trial variability 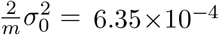 was larger than 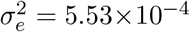, another incidence of a numerical singularity issue when estimating reliability through the TLM-based LME model in (11), likely a consequence when model assumptions were violated. Nevertheless, it is evident that the ICC estimate in this case was substantially attenuated.

Simulations were adopted to further explore different aspects of model performance for the condition-level contrast (Appendix E). The resulting patterns (Fig. 11) were largely similar to those from a single condition (Fig. 4). However, the difference between reliability estimates derived from the CLM-based method and those from the TLM-based approach was larger than for either of the two conditions or for the average between the two conditions. The magnitude of a contrast effect is much smaller than that of a condition or the average between conditions (see the mean column in Table 1). Thus, a contrast is associated with a larger variability ratio *R_v_*, which significantly attenuates the CLM-based estimate relative to the TLM-based estimate. As shown in Table 1, the ratio *R_v_* for the contrast is at least three times as large as that for single conditions.

### 2.5 Extension of the LME framework to BML

The extension from LME to BML is relatively straightforward. As apparent from numerical simulations and from applications to the Stroop data, the LME framework is ill-suited for reliability estimation: information on the precision and distribution of reliability estimates is not available, distribution assumptions (e.g., Gaussian) can be violated, sampling errors cannot be accommodated, and numerical failures occur. We now extend the LME formulations (5) and (11) to the BML framework. Specifically, we convert the two LME models with little modification for a single condition,

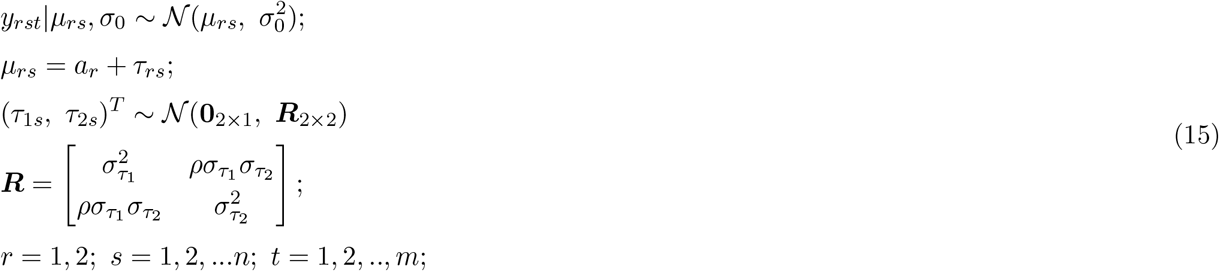

and a contrast between two conditions,

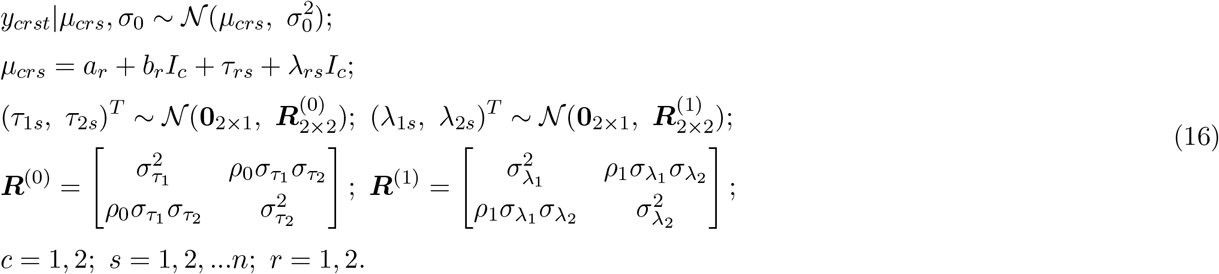

These BML models can be further extended to resolve the issues associated with the LME framework. For example, numerical instabilities and convergence issues would be largely avoided via modeling improvements including accommodating the data through various distributions such as Student’s *t*, exponentially modified Gaussian (exGaussian), log-normal, etc. More importantly, instead of providing point estimates without uncertainty information, the reliability estimation for *ρ* in (15), *ρ*_0_ and *ρ*_1_ in (16) can be expressed by their whole posterior distributions. Furthermore, sampling errors can be readily incorporated into the BML framework. For neuroimaging data, trial-level effects are usually not directly measured, but instead estimated through subject-level time series regression. Thus, it is desirable to include the standard errors of the trial-level effect estimates into the reliability formulation. With the hat notation for effect estimate 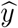 and its standard error 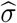, we broaden the two BML models (15) and (16), respectively, to,

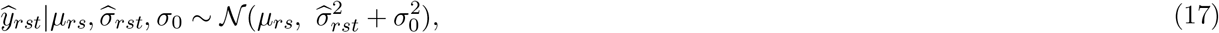

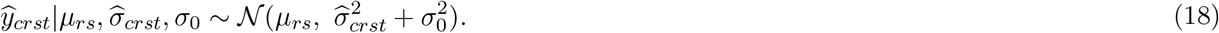

We illustrate the modeling capability and advantages of the Bayesian framework by directly comparing the models using the Stroop data. The LME framework assumes a Gaussian distribution as a prior and relies on its properties for convenient inferences through standard statistics such as Student’s *t*. However, this convenience comes at a cost: when the Gaussian prior is ill-suited, estimates can be inaccurate. Expanding on the work by Haines et al. (2020), who applied three likelihood distributions including Gaussian, log-normal and shifted log-normal to the data, we additionally considered Student’s *t* and exGaussian, because of their ability to handle skewed data and outliers. Moreover, instead of assuming a diagonal matrix ***R***^(0)^ for the variance-covariance structure of the varying intercepts *τ_rs_* as in Haines (2020), we adopted a generic ***R***^(0)^ for the BML model (16). In the Stroop dataset, the reliability estimation based on the Gaussian prior had the worst fit among the five priors (Fig. 5A-E), as was also reported by Haines et al. (2020). As exGaussian can accommodate skewed and outlying data such as reaction time which is lower-bounded, it outperformed all other prior distributions per the information criterion through leave-one-out cross-validation. This illustrates a principled approach to handling data distribution through modeling rather than data cleaning in common practice.

**Figure 5:**
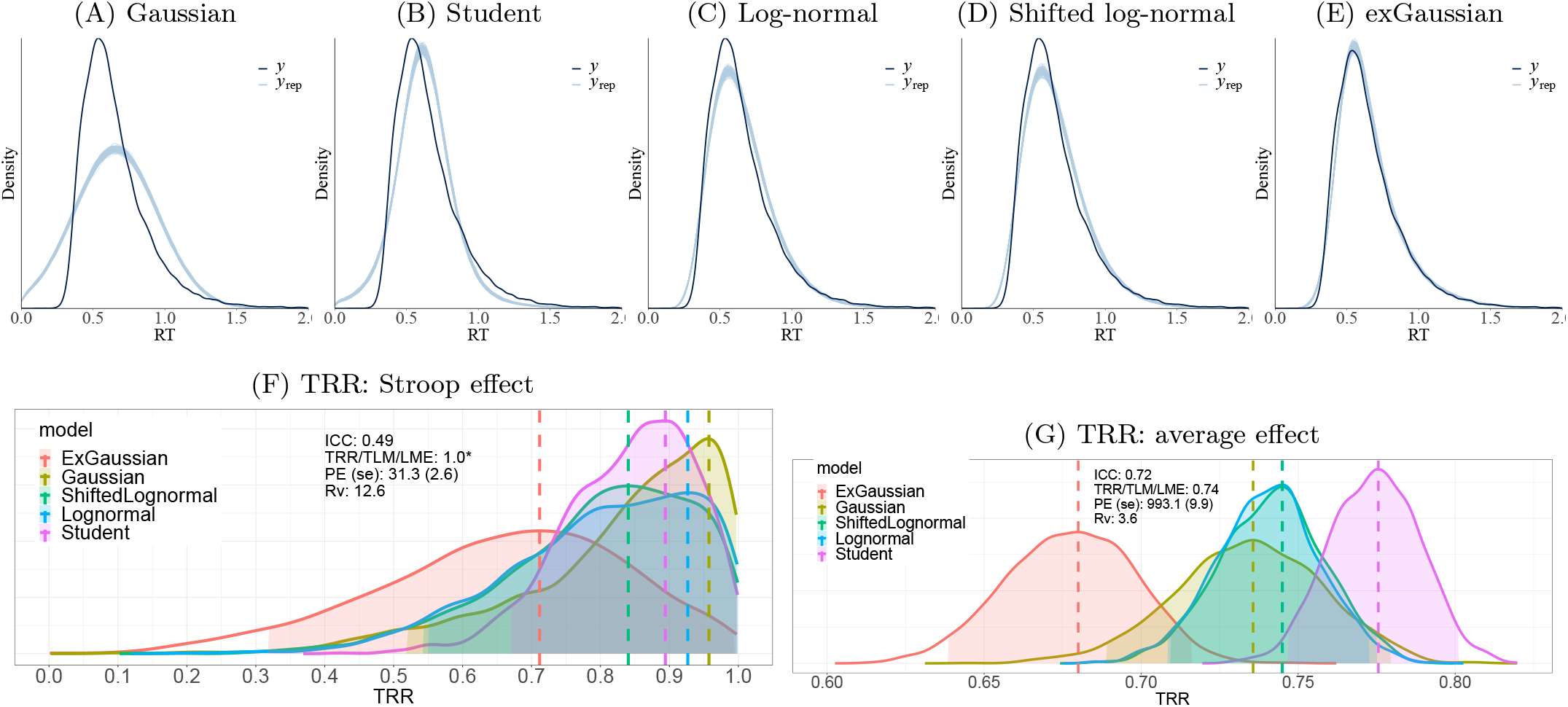
Model comparisons among five potential RT distributions for the Stroop dataset (Hedge et al., 2018). (A-E) Each of the five panels showing the posterior predictive density (light blue line) composed of 200 sub-curves, each of which corresponds to one draw from the posterior distribution. Overlaid is the raw data (solid black curves with linear interpolation). Overlaps between the solid black curve and the light blue line indicate fit of the respective model to the raw data. (F) Reliability distributions for the Stroop effect are shown for the five distributions. Reliability distributions are based on 2000 draws from MCMC simulations for each BML model. The dashed vertical line indicates the mode (peak) of each reliability distribution. Gaussian likelihood rendered the highest reliability with a mode of 0.96 but also exhibited the poorest fit to the data. The exGaussian distribution provided the lowest reliability with a mode of 0.71, while achieving the best fit among the five distributions. (G) Reliability distributions for the average between congruent and incongruent conditions are shown for the five distributions. Unsurprisingly, their uncertainty is much narrower than the contrast (F).

The rich information that can be derived from Bayesian modeling is worth noting. Unlike a simple point estimate under the conventional approach, we empirically construct the posterior distribution for a parameter through Monte Carlo simulations. The singularity problem (see the reliability estimate of 1.0 in Table 1) that we encountered with the LME model (11) is not an issue in the BML model (16). With a mode of 0.71 and a 95% highest density interval of (0.32, 0.99) for reliability estimate, it is clear that the ICC = 0.49 is substantially lower than the estimate derived through the BML.

We provide the program **TRR** for reliability estimation under BML. The two BML models, one for a single condition through (15) (or (17)) and the other for a contrast through (16) (or (18)), are implemented in the program **TRR** through Markov Chain Monte Carlo (MCMC) simulations using Stan (Carpenter et al., 2017) and the R package brms (Bürkner, 2017). Each Bayesian model is specified with a likelihood function, followed by priors for lower-level effects (e.g., trial, subject). The hyperpriors employed for model parameters (e.g., population-level effects, variances in prior distributions) are detailed in Appendix F. The program **TRR** is publicly available^2^ as part of the AFNI suite and can be used to estimate reliability for behavior and region-level neuroimaging data. Runtime ranges from minutes to hours depending on the amount of data.

## 3 BML modeling of reliability applied to a neuroimaging dataset

### 3.1 Data description

#### Modified Eriksen Flanker Task

Analyses in the current report used a subset of the subjects in Smith et al. (2020): 24 adults (>18 years; age: 26.81 ± 6.36) and 18 youth (<18 years; age: 14.01 ± 2.48). Subjects performed a modified Eriksen Flanker task (Eriksen and Eriksen, 1974) with 432 experimental trials during fMRI scanning in two separate sessions 53.5 ± 11.8 days apart. Participants were asked to identify, via button press, the direction of a central arrow, flanked by two arrows on either side. On half of the trials, the arrows were congruent with the center arrow (i.e., pointing in the same direction as the center arrow) and on the other half the arrows were incongruent with the center arrow (i.e., flanking arrows were pointing the opposite direction as the center arrow). The two trial types were randomized across the task with 108 additional fixation-only trials per session for a total of 540 trials per session. On each trial, a jittered fixation at a variable interval (300-600 ms) appeared on the screen followed by the Flanker arrows at a fixed time of 200 ms. The trial ended with a blank response screen of 1700 ms. The task was completed in four runs with three blocks per run to provide intermittent performance feedback to maximize commission errors. Stimulus presentation and jitter orders were optimized and pseudorandomized using the `make_random_timing.py` program in AFNI. Details regarding image acquisition and pre-processing are in Appendix G.

#### Subject-level Analysis

At the subject level, we analyzed the data with a time series model with regressors time-locked to stimulus onset reflecting trial type (incongruent, congruent) and error condition (correct, commission, omission). Regressors were created with a gamma variate for the hemodynamic response. The effects of interest at the condition level were two main contrasts: Cognitive Conflict (incongruent correct responses vs. congruent correct responses) and Error (incongruent commission errors vs. incongruent correct responses). All 42 participants were included in the conflict contrast, but only 27 participants had sufficient (≥ 20) commission errors in the incongruent condition to be included in the error contrast. For the conflict contrast, there were a total of 32005 trials across the two sessions of the task, which corresponds to 350 ± 36 incongruent trials and 412 ± 19 congruent trials across sessions per subject. For the subset of participants included into the error contrast, there were a total of 11366 trials available across both sessions, which corresponds to 331 ± 28 incongruent correct trials and 90 ± 27 incongruent commission errors per subject. We analyzed the subject-level fMRI data at the whole-brain level as well as at the region level using 12 Regions-Of-Interest (ROIs). We compare two reliability modeling approaches: a conventional CLM with regressors created at the condition level and TLM with trial-level regressors.

#### Region-of-interest (ROI) Selection

Seed coordinates for ROIs were selected using Neurosynth term-based meta-analyses with the terms “cognitive control” and “error”, the two main population-level effects of interest. Additionally, to derive ROIs outside the main population-level effects, we selected peak coordinates from term-based meta-analyses for the term “visual” and “default mode”. To select a reasonable number of peak coordinates, all four *z*-value maps (uniformity test for “cognitive control” and “error”, association test for the “visual” and “default mode” map), FDR-corrected to 0.01, were further thresholded to *z*-value of 10. Spheres with a 6-mm radius (57 voxels) were created for each of the 12 sets of peak coordinates derived from the surviving clusters. Six (6) spheres were derived from the “cognitive control” and “error” maps respectively, 4 spheres were derived from the “default mode” map and 2 spheres from the “visual” map. Spheres derived from the “default mode” and “visual” map were used for both conflict and error data for a total of 12 ROIs for each contrast.

### 3.2 Test-retest reliability estimation for behavioral data

Reliability estimates for the RT data of the Flanker task were relatively high.^3^ RT data were examined for the two main contrasts of interest: conflict (i.e., incongruent correct responses vs. congruent correct responses) and error (i.e., incongruent commission errors vs. incongruent correct responses). Trials with omissions were not considered. The RT values ranged within [4, 1669] and [9, 1686] ms for the conflict and error subset, respectively. As expected, the average RT values under each condition rendered relatively precise reliability estimates (Fig. 6B), and there was convergence between conventional ICC and BML for these estimates (conflict: ICC(3,1)=0.86, BML=0.86 (mode); error: ICC(3,1)=0.92, BML=0.90 (mode)). The reliability estimates for the RT contrast derived via the conventional ICC were 0.56 and 0.82 for conflict and error contrast, respectively. Under the BML framework, five distribution candidates were considered using the model (16): Gaussian, Student’s *t*, log-normal, shifted log-normal and exGaussian. Model comparisons and validations for each of the two RT contrasts (conflict and error) indicated that, similar to the RT data of the Stroop task (Fig. 5), the exGaussian distribution was the best fit alongside the Student’s *t*. Using the BML framework, the reliability for the conflict contrast, estimated as the mode of the reliability distribution, was higher than when estimated by the CLM-based approach, at a mode of 0.95. Some underestimation via the CLM is expected, given the associated variability ratio. There was less discrepancy for the error contrast between the two approaches with a mode of 0.88 through BML. However, for the error contrast, the BML model revealed that larger uncertainty was associated with the reliability estimation for this contrast (Fig. 6A).

**Figure 6:**
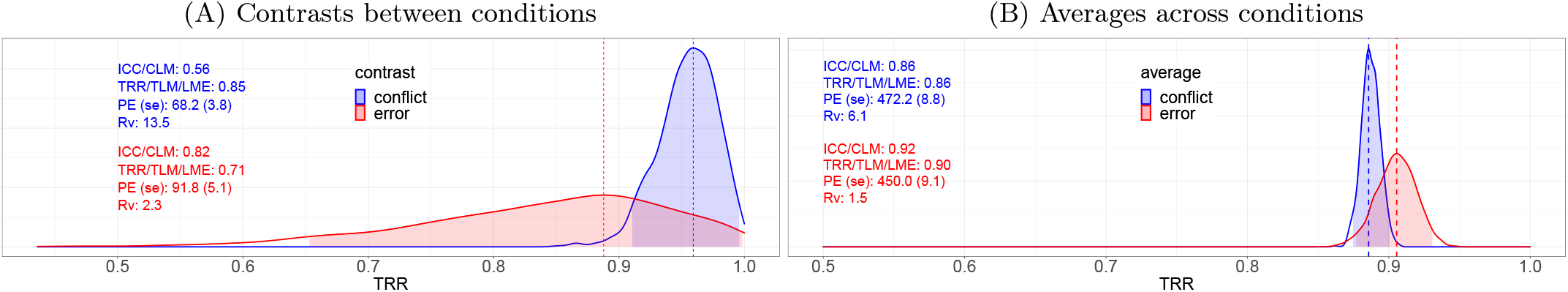
Reliability distributions of behavioral data (RT) from the Flanker task. The reliability estimates for the contrast (A) and average (B) between the two conditions are based on 2000 draws from MCMC simulations of the BML model with an exGaussian likelihood and are shown here as a kernel density estimate, which smooths the posterior samples. Each dashed vertical line indicates the mode (peak) of the perspective reliability distribution, and the shaded area indicates the 95% highest density interval. The reliability distribution for the conflict data was much more concentrated while the reliability for the error data was relatively diffuse. The magnitude of the variability ratio *R_v_* is a proxy to assess the degree to which the CLM-based method will generate a lower estimate. Population effects (PE) and standard errors (se) in milliseconds (ms) are shown for reference.

### 3.3 Whole-brain reliability estimation for neuroimaging data

As the BML model is not computationally feasible at the whole brain voxel-level, two modeling frameworks were adopted for whole brain analysis: ICCs using the program **3dICC** (Chen et al., 2018) with condition-level modeling (2) and the LME model (11) using the program **3dLMEr** (Chen et al., 2013) with trial-level modeling. Each model was applied to the average and the contrast between the two conditions for conflict (*n* = 42) and error data (*n* = 27) respectively, resulting in a total of eight analyses. Input for the ICC computation are condition-level contrasts, while for the LME computation trial-level conflict and error data was comprised of 32005 and 11366 three-dimensional volumes.

Individual differences were largely reliable among most voxels for the average between the two conditions as assessed via the CLM-based ICC (Fig. 7A,B). Similarly, reliability was in the moderate to high range for the average between the two conditions using the TLM-based LME (Fig. 7C,D). Both approaches rendered similar results for the average of conflict responses (Fig. 7, A vs C), while for the average of the error responses CLM-based estimates were noticeably lower, albeit not drastically (Fig. 7, B vs D). The divergence in estimates in the error contrast is likely due to a smaller number of trials for the error data, which is consistent with simulation results in Fig. 4B. Contrasts yielded lower estimates overall, with ICCs in the poor range (ICC(3,1) < 0.4, with the exception of adequate estimates in primary visual, parietal and motor regions). This is consistent with previous work (e.g., Elliott et al., 2020). ICCs were higher for the error than the conflict contrast in several regions. For contrasts, numerical failures occurred for most voxels in the TLM-based LME and thus results are not shown. Thus, in the next section, we explore the comparison between CLM/ICC and TLM using a region-level approach via the BML.

**Figure 7:**
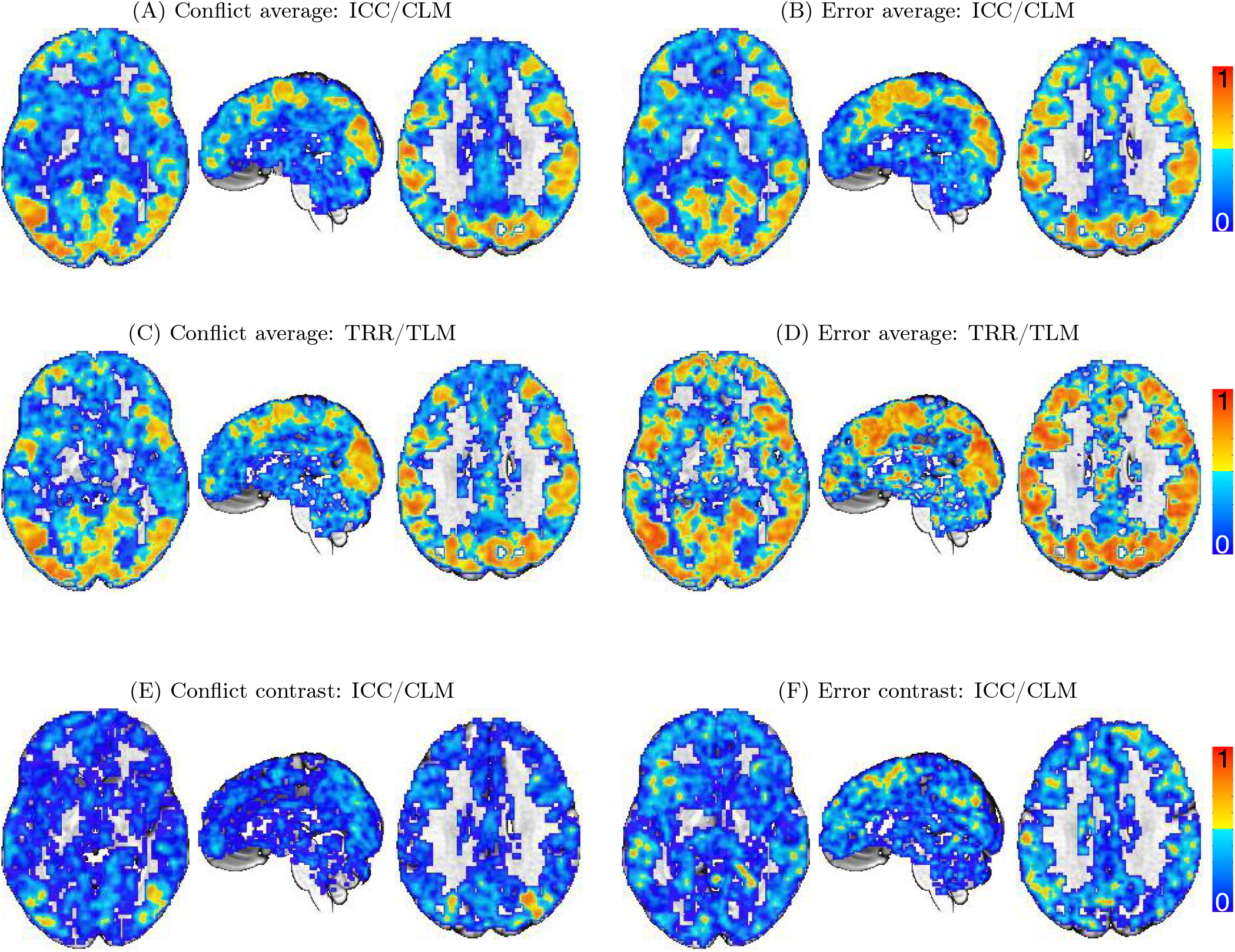
Whole-brain voxel-level reliability estimates for the fMRI Flanker data. The ICC values for the average conflict effect (A) showed negligible differences in reliability estimates between CLM and TLM/LME (C). In contrast, the ICC values for the average error effect (B) showed a moderate amount of underestimation compared to the reliability estimation (D) based on TLM/LME. The ICC values for the conflict (E) and error contrast (F) were much smaller than those for the average. Note that the TLM-based approach numerically failed for both contrasts at most voxels in the brain and the results are thus not shown. The three slices of axial (*Z* = 0), sagittal (*Y* = 14) and axial (*Z* = 28) planes are oriented in the neurological convention (right is right) in MNI space.

The relative magnitude of cross-trial variability in the Flanker data was substantial. Fig. 8A shows the distribution of the voxel-wise cross-trial variability across the brain, which has a mode of 20 and a 95% highest density interval (6, 86). These values are consistent with previous investigations of variability ratios in neuroimaging (Chen et al., 2020) and psychometric data (Rouder et al., 2019). In addition, the amount of trial-to-trial fluctuation can be appreciated in Fig. 8B, an illustration of the estimated effects at at the right middle temporal gyrus of a single subject.

**Figure 8:**
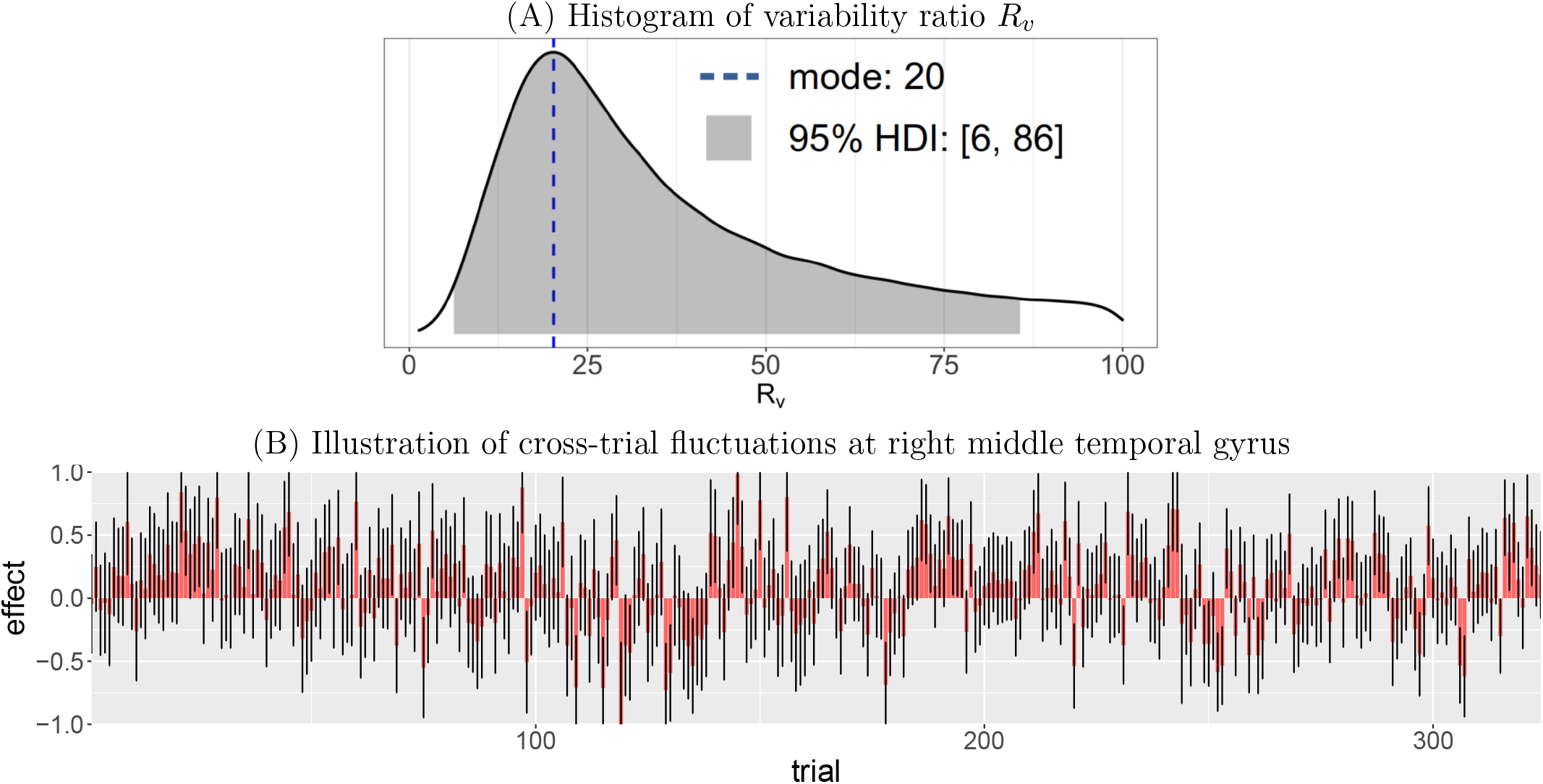
(A) Extent of cross-trial variability. The mode and 95% highest density interval (HDI) for the distribution of *R_v_* values in the brain are 20 and [6, 86], respectively. (B) The extent of variability across 324 correct responses to incongruent trials is illustrated at the right middle temporal gyrus of a subject. For each trial, the effect estimate is shown as an orange bar with a black segment indicating one standard error.

### 3.4 Region-based reliability estimation for neuroimaging data

Reliability estimation at the region level was performed through BML. The BML model (18) was adopted using the program **TRR** with the trial-level effect estimates from each subject as input. A Student’s *t*-distribution was utilized for cross-trial variability to account for potential outlying values (Chen et al., 2020). Standard errors of trial-level effects from the subject level were also incorporated into the BML model to improve robustness. The runtime was about 1.5 hours for each ROI through 4 Markov chains (CPUs) on a Linux computer (Fedora version 29) with AMD Opteron^®^ 6376 at 1.4 GHz.

Reliability estimates were in general higher for the average effect between the two conditions than their contrast. Most regions exhibited adequate to excellent reliability for the average effect between the two conditions for both conflict and error data with a few exceptions (R precuneus and R angular gyrus). However, there was some variability in the precision of these reliability estimates (Fig. 9). For the contrast, we found large variations across regions in terms of both magnitude of the reliability estimate and precision (Fig. 10). Some regions showed moderate to high reliability with relatively high precision, some regions exhibited high reliability with a wide range of uncertainty, others were difficult to assess as their reliability distributions were very diffuse. For regions with such substantial amount of uncertainty, an estimate through the ICC would be largely meaningless. The following regions demonstrated high reliability with relatively high precision: left MOG and right MTG for the conflict contrast (Fig. 10A); left SMA and PreCG for the error contrast (Fig. 10B). Some regions had reasonably high reliability but with moderate to poor precision (e.g., left SMA, left IPL, right angular gyrus for conflict contrast; left IL, right IFG, left and right IFG, MOG and right MTG for error contrast). As demonstrated above, the variability ratio can be used as a proxy for the degree of underestimation via CLM, and provides an explanation as to why such heterogeneity exists. Ten (10) out of the 12 ROIs had a substantial range of uncertainty associated with their reliability estimate for both conflict and error contrasts (Fig. 10). These estimates were for the most part associated with a large variability ratio. It is for this reason that we advocate that reliability should not simply be expressed as a point estimates through the conventional ICC.

**Figure 9:**
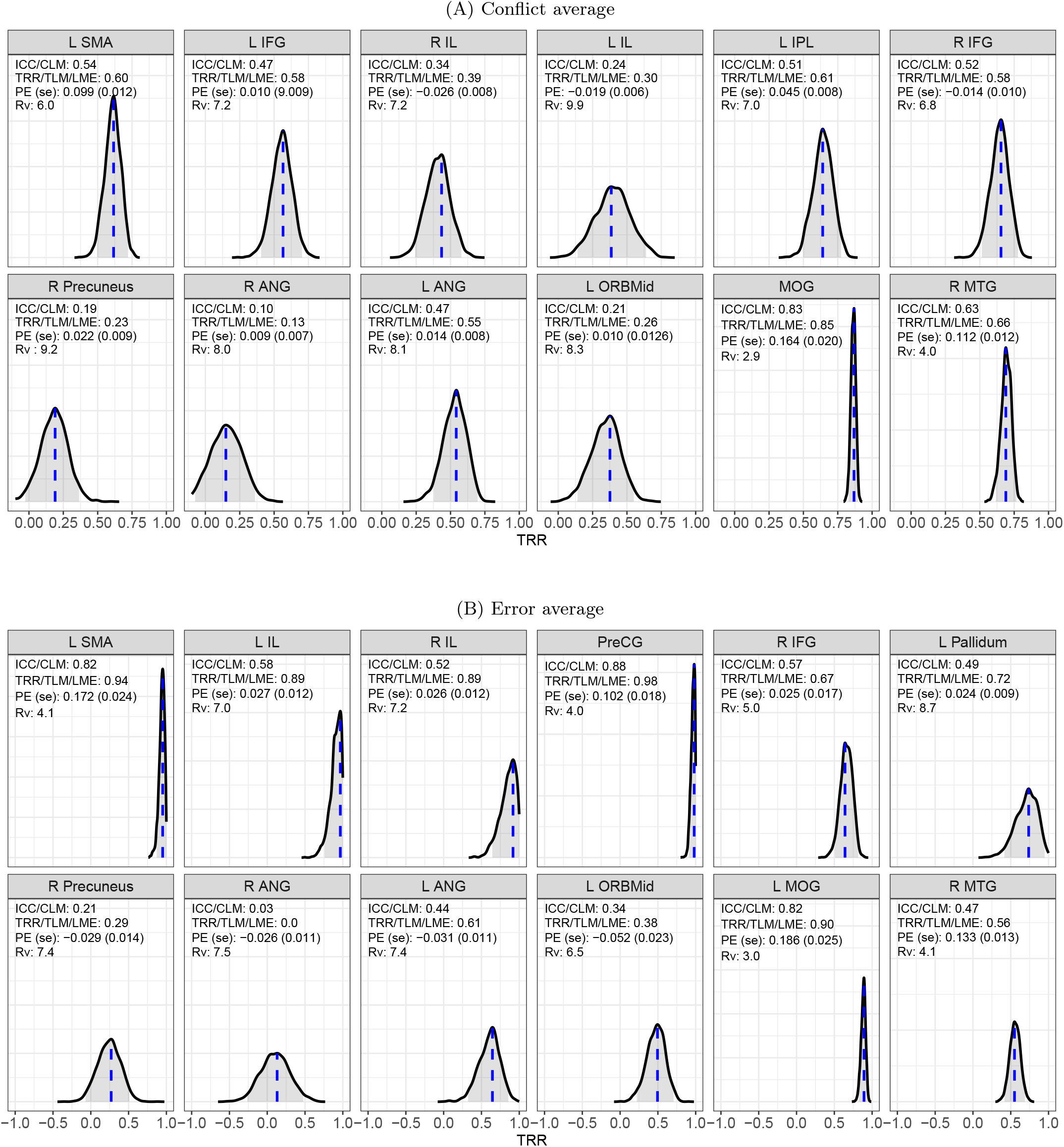
Reliability distributions for the average effect between the two conditions at 12 regions. The reliability estimates for the average effects of conflict (A) and error (B) were obtained using the program **TRR** based on 2000 draws from MCMC simulations of the BML model with a Student’s *t*-distribution. Each blue vertical line indicates the mode (peak) of the reliability distribution, and the shaded area shows the 95% highest density interval. Four quantities are listed in the density plot for each region: conventional ICC based on CLM, reliability estimated through TLM, population effect (PE) estimate plus its standard error (se), and variability ratio *R_v_*. Asterisk (*) indicates numerical problems (either singularity or convergence failure) under LME. Abbreviations: R: right, L: left, SMA: supplementary motor area, IFG: inferior frontal gyrus, IL: insula lobe, IPL: inferior parietal lobule, PreCG: precentral gyrus, MOG: middle occipital gyrus, MTG: middle temporal gyrus, ANG: angular gyrus, ORBmid: middle orbital gyrus.

**Figure 10:**
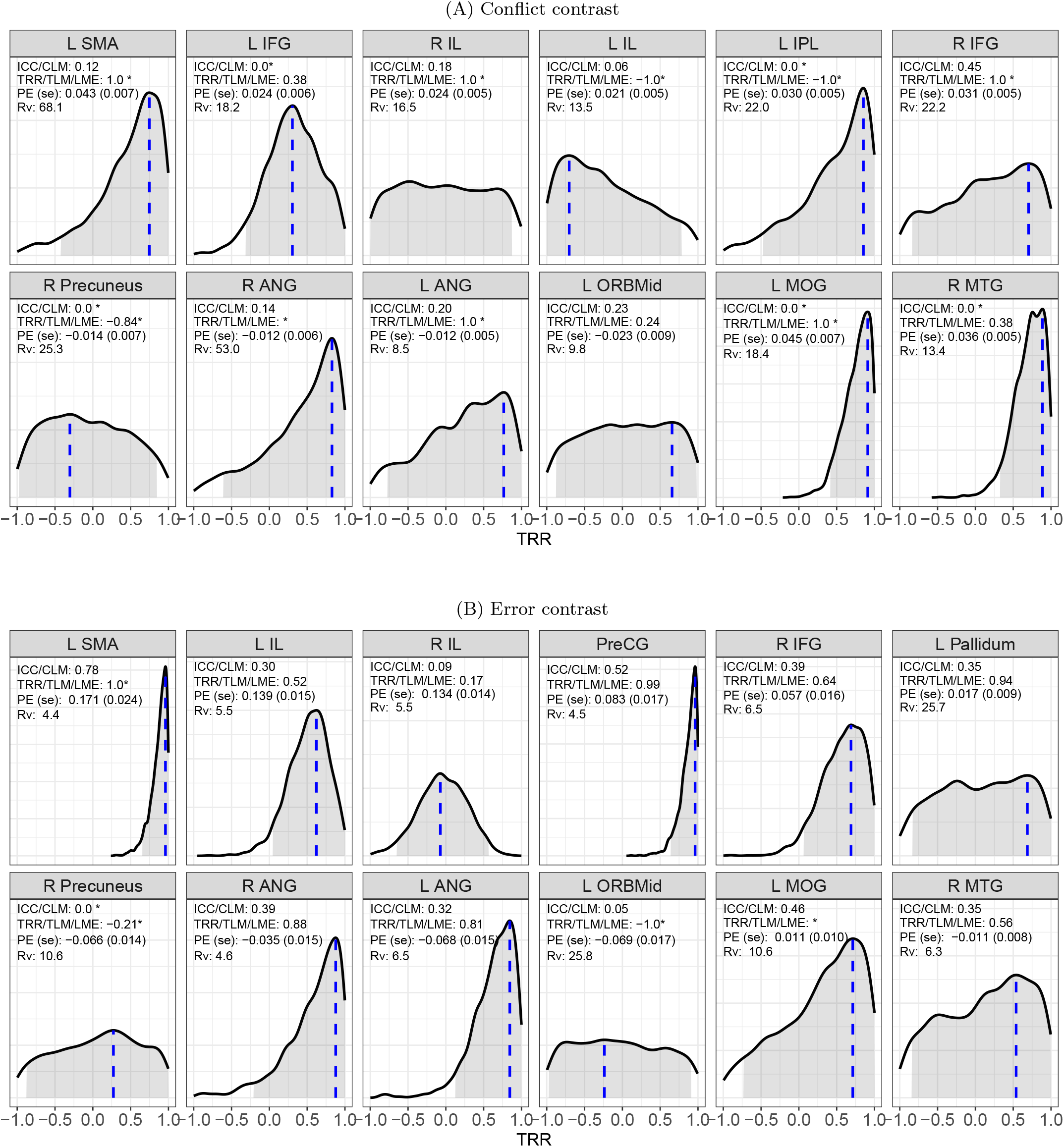
Reliability distributions for the two contrasts of the Flanker task per ROIs. The reliability estimates for the conflict (A) and error (B) contrast were obtained using the program **TRR** based on 2000 draws from MCMC simulations of the BML model with a Student’s *t*-distribution. TEach blue vertical line indicates the mode (peak) of the respective reliability distribution, and the shaded area shows the 95% highest density interval. Four quantities are listed in the density plot for each region: conventional ICC based on CLM, reliability estimated through TLM, population effect (PE) estimate plus standard error (se), and variability ratio *R_v_*. Asterisk (*) indicates numerical problems (either singularity or convergence failure) under LME. Abbreviations: R: right, L: left, SMA: supplementary motor area, IFG: inferior frontal gyrus, IL: insula lobe, IPL: inferior parietal lobule, PreCG: precentral gyrus, MOG: middle occipital gyrus, MTG: middle temporal gyrus, ANG: angular gyrus, ORBmid: middle orbital gyrus.

### 3.5 Implications for practice

The advancements through the BML are substantial for any researcher hoping to assess reliability for their task-based imaging data. It is specifically the additional precision information that is crucial. The current version of the Flanker task was selected as a task that, with its two contrasts, covered several scenarios in the literature: The conflict contrast, with its 432 experimental trials per session likely represents the upper range of trial-level repetitions. The error contrast with a minimum of 20 trials in the incongruent commission error condition has just barely enough trials to derive robust estimates in a conventional analysis. Additionally, the number of participants for the conflict contrast (42) was almost double of that of the error contrast (27). The variability ratio *R_v_* was generally larger for the conflict than the error contrast, both in the behavioral and imaging data (albeit variable across regions). Half of the trials in both contrasts overlap (i.e., incongruent correct responses are used across both contrasts). Hence, commission error responses must be driving the increased reliability in the error contrast compared to the congruent correct trials included in the conflict contrast. In other words, despite relatively few instantiations in the task, commission error responses may generate a more consistent signal within person that distinctly differs between individuals. Alternatively, it may be that, despite involving the same visual display, the correlation between conditions is smaller in the error contrast. The discrepancy in the psychological processes engaged during the error contrast is more pronounced compared to the conflict contrast, where differences in cognitive demands are more subtle. Interestingly, the error contrast of RT showed large uncertainty around reliability estimates compared to the conflict contrast, which had a more narrow reliability distribution. No such visible differences in uncertainty emerged between the two contrasts in the region-level analysis of the imaging data. It is plausible that while behaviorally errors are few and inconsistent, generated via different mechanisms (i.e., short error RTs due to premature, anticipatory responses or flanking distractors, long error RTs due to attention lapses), these errors result in a common, intra-individually relatively consistent but inter-individually distinct error signal in the brain. This tentative explanation highlights the need to more systematically use BML to explore reliability across different cognitive processes and experimental designs. Most importantly, our initial results provide a more nuanced assessment of task fMRI reliability than recent reports: reliability variability is large, even within the same contrast across regions, both in magnitude and precision of reliability estimates; understanding the source of trial-level variability may provide an important avenue to improve reliability in future work.

We note that for TLM-derived estimates to be most useful, it would be important to also apply a comparable TLM approach to the corresponding subject-level analysis (Chen et al., 2020). It is essential to assess reliability of the exact measurement that is used in the experiment; reliability estimates are informative upper bounds for expected correlations between two measures (Spearman, 1904). Hence, if subject-level analysis is adopted to obtain CLM-based reliability estimates for the conventional ICC computation, while less precise, may still represent a useful indicator of reliability. Caution, however, is advised – our investigation has provided evidence that there are several scenarios in which CLM-based point estimates would be misleading.

## 4 Discussion

For almost all scientific endeavors, it is critical that measurements taken are of adequate temporal stability. Here, we adopt a new modeling framework for reliability estimation in behavioral and imaging tasks that (a) reflects the data hierarchy including individual trials, (b) handles outliers and skewness in a principled way, and (c) properly quantifies uncertainty. Our investigation illustrates the benefits of constructing an adaptive model that preserves the hierarchical integrity of the data and the cost of data reduction. Several key findings emerged in the step-by-step process by which we built the reliability model formulation: (a) reliability is conceptually distinct from population effects, (b) the conventional CLM-based approach yields lower reliability estimates in cases where condition-level effects are assessed through an ensemble of trials and the trial-level variability is large relative to between-subject variability, (c) trial-level variability is surprisingly large in both behavioral and neuroimaging data, and (d) a large number of trials is important to adequately assess reliability. We will discuss each of these in turn.

### 4.1 The relationship between population-level effects and test-retest reliability

We demonstrate that population effects are not necessarily tied to the reliability of individual differences. As illustrated in Fig. 1, all four possible scenarios can occur: large population effects can have strong or weak reliability, and the absence of population-level effects does not preclude high reliability. Researchers usually choose to examine reliability of neural measures in regions that exhibit strong population effects; strong evidence of population-level effects may pique the researcher’s interest in exploring the reliability of individual differences. However, limiting the search space to population-level effects may be too narrow.

The common practice of dichotomizing results into “significant” and “not significant” via multiple testing adjustment specifically exacerbates this problem (Chen et al., 2021a). This can mean missed opportunities to estimate the reliability for regions involved in the task or regions that, while not showing involvement “on average”, are relied upon by a subset of individuals. For example, the statistical evidence associated with the four regions of the Flanker dataset (R Precuneus, R ANG, L ANG and L ORBMid) was not strong enough per the currently adopted criteria for multiple testing adjustment, and would not have been part of the current reliability exploration if the ROI selection had been solely based on the statistical strength at the population level. It is important to note that to maximize the utility of estimates derived from the hierarchical model, it would be crucial to match the analytic approach with a proper trial-level modeling at the subject level. Previous work (Chen et al., 2020) has detailed how to adopt trial-level modeling.

### 4.2 BML as an adaptive solution for reliability estimation

Recent reports using the ICC metric suggest that the reliability of commonly used fMRI tasks is inadequate. Issues of reliability are now front and center in the field, and have motivated new modeling approaches for behavioral and imaging data with multivariate approaches recently highlighted as an avenue with more promising test-retest reliability (e.g., Noble et al., 2021). Here we show that, through theoretical reasoning and effect partitioning, the hierarchical modeling structure closely characterizes trial-level variability and improves the estimation accuracy of test-retest reliability relative to the conventional ICC formulation (e.g., formulas (6) and (12)). In addition, the robustness of TLM-based reliability estimation was verified in simulations through the successful retrieval of ground truth. In contrast, the conventional ICC formulation through CLM did not correctly render the simulated reliability; furthermore, the underestimation of reliability via the conventional ICC is clearly illustrated (Figs. 4 and 11) and was also demonstrated using experimental data (Figs. 9 and 10). Furthermore, the attenuation by the conventional CLM-based approach is linearly associated with the magnitude of reliability, with the extent of underestimation depending on two factors: the trial sample size and the relative magnitude of cross-trial to cross-subject variability. The larger the relative cross-trial variability, the more severe the underestimation. Therefore, a reliability estimate, when implicitly “contaminated” by cross-trial variability, is sensitive to trial sample size. As a result, the reliability estimates reported in the literature, even on the same task, may not necessarily be comparable across experiments, leading to a portability problem (i.e., the independence of a statistical metric on sample sizes, Rouder and Haaf, 2019). A recent study by Han et al. (2021) corroborated the dependence of the CLM-based ICC on the number of trials; the authors report test-retest reliability values of 0.84, 0.74, and 0.46 for an experiment with 70, 30, and 5 trials, respectively.

Hierarchical models such as LME and BML through TLM allows the researcher to disentangle trial-level effects from other sources of variance, arguably providing a more accurate assessment of reliability. In other words, even though trial-level effects are of no interest, accounting for them in the model separates the cross-trial variability from the cross-subject counterpart and allows the accurate estimation of the correlation structure across repetitions. With a data structure involving five levels (Fig. 3), a hierarchical model closely follows the underlying data generative process and simultaneously incorporates all the information available through proper regularization. As a result, one effectively gains high predictive accuracy for reliability estimation. On the other hand, one could continue to adopt the conventional ICC formulation with the condition-level effect estimates from each subject. However, it is worth pointing out that ignoring the cross-trial variability does not mean the exclusion of the variability’s presence in the ICC computation. Rather, as our theoretical discourse and simulations show here, an attenuated estimate of test-retest reliability will occur.

The BML framework is well-suited for reliability estimation. The adoption of the Bayesian formulation is mostly not intended to inject prior information, but to overcome several limitations of the LME approach through Monte Carlo simulations. Despite their accommodation of multilevel data structure, LME models can encounter numerical difficulties. In addition, LME models only provide point estimates with no easy access to uncertainty and have limited flexibility in handling deviations from a Gaussian distribution (such as data skewness and outliers). BML can overcome theses LME-associated issues. For example, rather than censoring data through arbitrary thresholding, BML uses a principled approach and can adapt to the data through a wide variety of distributions (see Fig. 5). The information contained in posterior distributions is another benefit, showing subtle differences in distributions and allowing for straightforward interpretations. Despite a large amount of noise unaccounted for in fMRI data and a high variability ratio, the modeling investment is worthwhile and reveals reliability values above 0.9 at some regions with our neuroimaging data (Figs. 9,10).

Model comparison and validation are important in improving the accuracy of reliability estimation. We note that a Gaussian assumption adopted usually as default may not always be an appropriate choice. The choice of assumption reflects prior knowledge about the specific data generating mechanism (i.e., bounded, skewed) and realities about variability in measurement (e.g., outliers). Hence, we recommend that, unlike in the conventional framework where this choice is largely epistemological, any assumption adopted as a prior or likelihood should be calibrated by the data and further cross-validated though, for instance, posterior predictive checks. Even for data with substantially less complexity than imaging data, such as RT, a consensus has not been reached in terms of what distribution is a “default” best fit, although most investigations have leaned towards adopting either an exGaussian or a shifted log-normal distribution. This is because even for this ‘simple’ measure, complexity arises through mechanisms such as linear increases of uncertainty with increasing means (Wagenmakers and Brown, 2007).

### 4.3 Two crucial factors in reliability estimation: cross-trial variability and trial sample size

Large cross-trial variability is the main cause for lower estimates from CLM, numerical failures in LME modeling and poor precision of reliability estimates. Only in scenarios where cross-trial variability is roughly the same order of magnitude as or smaller than cross-subject variability, the conventional ICC through CLM and the reliability estimates based on TLM will yield similar results. However, experimental data from both behavioral and neuroimaging investigations point to a much larger cross-trial variability than cross-subject variability in most tasks. Specifically, the variability ratio *R_v_* may reach up to 10 for simple effects and go beyond 20 for contrasts. Trial-level effects fluctuate substantially; these fluctuations do not appear to have a clear pattern (Chen et al., 2020) and cannot be accounted for by habituation, fatigue or sensitization. At the same time, these trial-level fluctuations are not purely random: there is a high degree of bilateral synchronization within the same subject (Chen et al., 2020). Some of this variance may be accounted for by behavioral measures such as reaction time (i.e., momentary lapses in attention) or more systematic stimulus ratings that can be modeled through trial-level modulation analysis at the subject level. It is also plausible that brain regions may constantly undergo some intrinsic fluctuations – external stimuli or tasks may be surprisingly small constraints stacked on top of large intrinsic neuronal activities. Strong evidence based on electroencephalography indicates that the substantial cross-trial variability is mainly caused by the impacts of ongoing dynamics spilling over from the prestimulus period that dwarf the influence of the trial itself (Wolff et al., 2021). Additionally, cross-trial variability might stem from suboptimal modeling in subject-level time series regression due to (i) a substantial amount of confounding effects not properly accounted for and/or (ii) poor characterization of varying hemodynamic response across trials, conditions, regions and subjects through a fixed basis function. However, large cross-trial variability also exists in behavioral tasks although to a slightly lesser extent (Rouder et al., 2019). Thus, suboptimal modeling likely is not the main (or only) culprit; understanding the origins of this variability remains an important future endeavor.

The number of trials plays a crucial role in achieving a reasonably precise reliability estimates. We illustrated the strong relationship between accuracy of the CLM-based and trial sample size. It has been generally assumed that a larger subject sample helps achieve high statistical efficiency (i.e., small standard error) of test-retest reliability (Shoukri et al., 2004). However, while subject sample size plays a role in stability and precision of reliability estimates, trial number has a much stronger impact on reliability estimates and efficiency, as shown in the formulas (6) and (12) as well as simulation results in Figs. 4 and 11. The central limit theorem is pivotal to many modeling frameworks including the conventional CLM. However, the asymptotic property of both unbiasedness and Gaussianity relies on large sample sizes, an assumption that will not necessarily be met in practice. Ensuring a large enough sample size, especially in terms of trials, for reliability estimation is an important issue that has received far too little attention. Based on empirical data, we showed that the trial-to-trial fluctuations are at least a few times larger than cross-subject fluctuations (Fig. 8), violating the underlying assumption of the CLM-based approach. Thus, trial sample sizes in typical experimental designs are usually not large enough to estimate condition-level effects with little uncertainty. Without the ability to accommodate cross-trial fluctuations, the conventional CLM-based estimation of reliability of individual differences leads to substantially lower estimates. Of note, the issues surrounding the large trial-level variability also apply to conventional population-level analysis and hierarchical modeling strategies can similarly be applied (Chen et al., 2020; Chen et al., 2021b).

Because cross-trial variability is often high, a substantial number of trials may be required to achieve a reasonable precision of estimates. As shown in our investigation, the uncertainty of reliability estimates, as represented by their standard error or highest density interval, often remains wide when cross-trial variability is large. Uncertainty depends on four factors (Figs. 4 and 11): reliability magnitude, variability ratio, trial and, to a lesser extent, subject sample size. Among the four factors, only the sample sizes can be easily manipulated. Also, as shown in experimental results (Figs. 5, 6, 9 and 10), the precision of reliability estimates varies substantially across brain regions and between single conditions and contrasts. To dissolve those diffuse posterior distributions of reliability estimates, a few hundred or even more trials may have to be adopted. In practice, such experimental designs may not be feasible due to their time burden on the subject and financial burden on the experimenter.

### 4.4 Difficulty of obtaining high reliability precision for a contrast

For effects of a single condition or the average among two or more conditions, it is relatively easy to achieve reasonable reliability precision. Empirical data indicate that cross-trial variability *σ*_0_ is larger than cross-subject variability *σ_τ_r__* with a variability ratio *R_v_* less than 10. Thus, it is possible, with a sizeable trial sample size (possibly larger than what is typically adopted in the field), to obtain reliability estimates within a small or moderate amount of uncertainty (Figs. 5, 6 and 9). Consequentially, one may be able to estimate reliability under a linear mixed-effects framework through trial-level modeling (Fig. 7C,D).

In comparison, a reasonably high precision of reliability estimation for a contrast might be harder to obtain. Cross-trial variability *σ*_0_ measures the trial-level fluctuations per condition (and per subject) for a single condition or a contrast. However, when a contrast is of interest, cross-subject variability *σ_τ_* measures the fluctuations relative to the contrast, as characterized in the parameters *λ_rs_* in formulations (11) and (16). The contrast between two conditions is usually a few times smaller in magnitude. For example, suppose that the magnitude of the BOLD response in the congruent and incongruent condition is 1.0% and 0.8% at a brain region, respectively. Their contrast of 0.2% would be 4-5 times smaller than the magnitude of each condition alone. Yet, cross-trial variability *σ*_0_ remains roughly the same regardless of the effect (i.e., contrast, a single condition or cross-condition average). Thus, the relative magnitude of cross-trial variability would be much larger for the contrast than a single condition, leading to a sizeable variability ratio *R_v_*. In other words, the cross-subject effects *λ_rs_* often will get dwarfed by cross-trial variability *σ*_0_, resulting in a large uncertainty for the reliability estimation of a contrast. The variability ratio *R_v_* can differ substantially across regions and may be so large in some regions that the requirement for trial sample sizes may become practically unfeasible. Nevertheless, as is evident from Fig. 10, some brain regions still achieve high reliability estimates with reasonable precision. Even though it may be difficult to achieve a high precision for reliability estimates, the resulting posteriors from a BML model would likely still encapsulate the distribution shape regardless of its centrality or diffusivity. Overall, we recommend the adoption of the BML framework for its flexibility to closely characterize the data structure.

### 4.5 Three types of generalizability

Scientific investigations aim to gain knowledge through legitimate generalization. From limited samples and properly built models, researchers should be able to draw broader conclusions that extend far beyond specific experiments. This generalization is made possible through inferences regarding a hypothetical population based on the data at hand. Three types of generalizability are relevant in the current context: sample average to population average, stability of sample individual differences across time to test-retest reliability, and individual trials (stimuli) of a condition to a stimulus category.

Population-level effects generalize through measurement units of subjects and trials. They are usually conceptualized at the top of the data hierarchy and are modeled as fixed effects under the conventional LME framework. Therefore, cross-subject and cross-trial effects are considered random fluctuations. In contrast, test-retest reliability concerns a different type of generalizability: the consistency of individual differences. Unlike population-level effects, which are denoted as “fixed” in a statistical model, reliability is characterized by subject-level effects that vary across subjects and are considered “random” under the LME framework. Cross-subject fluctuations are expected to be consistent and systematic – hence, generalization pertains to the stability of between-subject variance (relative variations around the population effects) across time. Due to their smaller effect sizes compared to their population effect counterparts, subject-level effects and reliability are often more subtle.

Lastly, generalization to a stimulus category (e.g., faces) from individual examples happens through the units of trials. To generalize from a sample of trails, these need to be explicitly modeled. For example, under the conventional linear mixed-effects (or hierarchical) framework, the trial-level effects are explicitly characterized as random effects in the model. On the other hand, condition-level modeling, where trial variance is not modeled but collapsed across, does technically not allow for generalization to a stimulus category because it assumes no cross-trial variability (which does not reflect reality). Fig. 2f-g illustrates that generalization from individual stimuli (i.e., trials) to a stimulus category is only possible if these are represented in the model.

### 4.6 Limitations

BML modeling remains practically limited to region-level reliability estimation. Currently, it is not possible to apply the BML framework at the whole-brain voxle-wise level. Because long chains of iterations are required to obtain stable numerical simulations under the Bayesian framework, the computational cost of BML is usually high for large datasets. Thus, its application is currently limited to behavioral and region-based neuroimaging data.

Trial-level modeling requires careful experimental designs. When each trial is modeled separately at the subject level, the risk of high correlations or multicollinearity may arise among the regressors. To avoid such potential issues, trial sequence and timing can be randomized to reduce multicollinearity using tools such as `make_random_timing.py` in AFNI or `optseq`^4^. However, even if statistically separable, trial-level effect estimates can be unreliable and estimation procedures are largely limited to the common approach of assuming a fixed-shape hemodynamic response for most experimental designs. As a substantial amount of variability exists across tasks, brain regions, subjects and even trials, fixed shapes might misidentify trial-level effect magnitudes, resulting in compromised reliability estimation.

## 5 Conclusion

The conventional CLM-based approach, when adopted for datasets with many trials comprising a condition of interest, will lead to attenuated test-retest reliability estimates compared to the proposed hierarchical model that adequately accounts for trial-level variability. The TLM-based approach provides two concrete advantages relative to the condition-level approach: 1) it precisely partitions the hierarchical structure of the data down to individual trial effects, leading to more precise estimates of subject-associated variability, 2) through a Bayesian approach, the BML model allows the experimenter to accurately estimate the test-retest reliability and its uncertainty, which are critical for proper interpretation. We offer two programs, available to the public, **TRR** and **3dLMEr**, to apply these procedures. In addition, we suggest that reliability should be reported with either a full posterior distribution or a mode combined with its highest density interval. A large number of trials might be required to generate reliability estimates with acceptable precision (i.e., relatively low uncertainty) especially for effects of smaller magnitude such as contrasts between two conditions.

## Acknowledgments

The research and writing of the paper were supported (GC and RWC) by the NIMH and NINDS Intramural Research Programs (ZICMH002888) of the NIH/HHS, USA. Data collection was supported (DSP) by the NIMH Intramural Research Program (ZIAMH002781). Our work was inspired by the modeling platforms of Haines et al. (2020) and Rouder and Haaf (2019). We are appreciative of the technical support from the `Stan` (Carpenter et al., 2017) and R (R Core Team, 2019) communities. Most of the modeling work was performed in `Stan` through the R packages `brms` (Bürkner, 2018) and `Ime4` (Bates et al., 2015). The figures were generated with the R package `ggplot2` (Wickham, 2009). This work utilized the computational resources of the NIH HPC Biowulf cluster (http://hpc.nih.gov). We thank the anonymous reviewers for their critical reading with thoughtful suggestions that helped improve, clarify and contextualize our manuscript.

## Appendices

## A Attenuated reliability estimation via the CLM-based method for a single condition

We seek to derive the reliability under the LME framework through TLM. It is worth noting that, under the LME formulation (5), ICC can be conceptualized as the correlation between the two cross-trial averages at the subject level 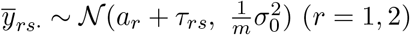 with the homoscedastity assumption between the two repetitions *σ*_*τ*1_ = *σ*_*τ*2_ = *σ*_*τ*_. With the notations

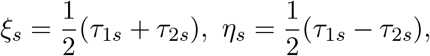

we have

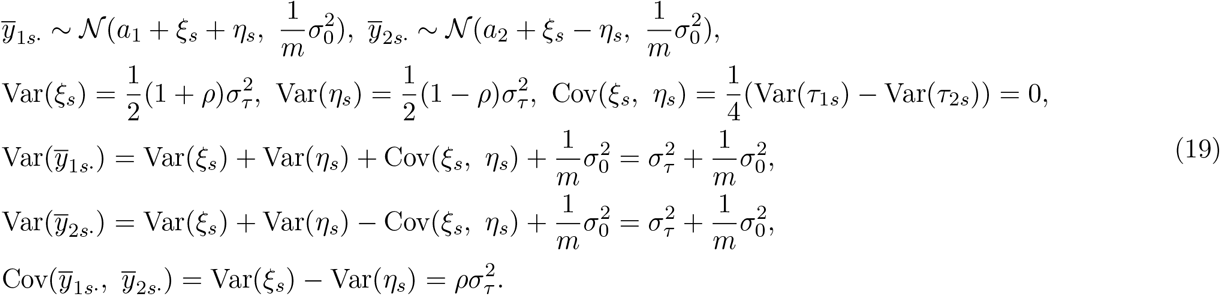

Through the notations

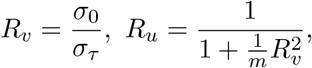

it becomes clear that ICC can be directly expressed as the function of *ρ*

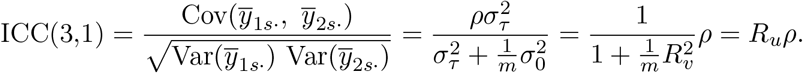

The variability ratio *R_v_* characterizes the magnitude of cross-trial variability *σ*_0_ relative to the cross-subject variability *σ_τ_*, and the parameter *R_u_* encapsulates the rate of underestimation by CLM. It is quite revealing that the extent to which estimates are lower depends on two factors, the trial sample size *m* and the relative magnitude of cross-trial variability *R_v_*.

The underestimation by CLM can be conceptually corrected. Under the homoscedasticity assumption, the derivations (19) indicate that 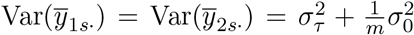. That is, the variability of cross-trial averages is composed of two components, one associated with cross-subject variability *σ_τ_* and the other with cross-trial variability *σ*_0_. Thus, if the cross-trial variability *σ*_0_ is known, we could restore the accuracy of ICC by removing the trial-related variance component from the denominator of the ICC formulation (3),

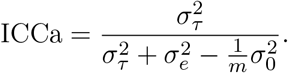

## B Simulations with a single condition

Simulations were conducted for reliability with a single condition. Below are the manipulation parameters for the two models of condition-level (2) and trial-level modeling (5) with two repetitions of data collection:

1. homoscedasticity with fixed scaling parameters across sessions: *σ*_*τ*_1__ = *σ*_*τ*_2__ = *σ_τ_* = 1,
2. 4 different reliability values: *ρ* = 0.3, 0.5, 0.7 and 0.9,
3. 4 different sample sizes of subjects: *n* = 20, 40, 100 and 200,
4. 4 different sample sizes of trials per repetition: *m* = 20, 40, 100 and 200,
5. 4 different ratios of cross-trial relative to cross-subject variability: 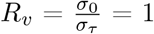 and 10,
6. 2 different sets of population-level effects across the two repetitions: (*a*_1_, *a*_2_) = (0, 0) and (1.0, 0.9),
7. 3 different approaches to assessing cross-session reliability:

a. conventional ICC estimated with ANOVA/LME (2) through aggregation across trials,^5^
b. reliability *ρ* estimated through LME (5),
c. conventional ICC adjusted by removing the cross-trial variability 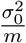

Each of these 4 × 4 × 4 × 4 × 2 × 3 = 1536 combinations was simulated 1000 times, using the function *lmer* in the *R* package `Ime4` (Bates et al., 2015) with the following iterative steps.

i. For each subject *s*, obtain subject-level effects during the two repetitions through random sampling:

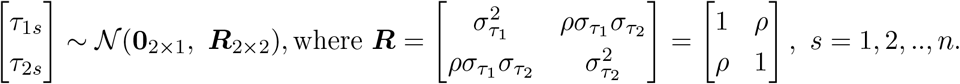
ii. Draw the simulated data under LME (5):

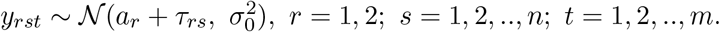
iii. Solve the two LME models of (2) and (5).
iv. Recover the simulated parameters including the three reliability estimates. Specifically, the conventional ICC is obtained through the formula (3) while the reliability is estimated through *ρ* in (5). In addition, the adjusted ICC is obtained by removing the cross-trial variability 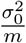 through the formula (8).

## C Correlation structure among the varying intercepts and varying slopes in the LME model (11)

With two conditions and a 2 × 2 factorial structure, we denote *μ_crs_* as the *s*-th subject’s condition-level effects (*c* = 1,2; *r* = 1,2; *s* = 1,2,…, *n*) and assume that the four effects associated with each subject follow a quad-variate Gaussian distribution,

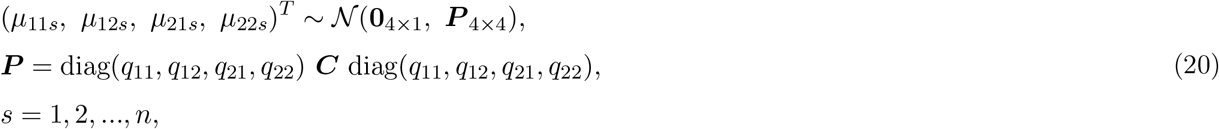

where ***P*** and ***C*** are the variance-covariance and correlation matrices for the four effects. With the symmetry assumptions corr(*μ*_*c*1*s*_, *μ*_*c*2*s*_) = *π* (*c* = 1, 2), corr(*μ*_1*rs*_, *μ*_2*rs*_) = *θ* (*r* = 1, 2) and corr(*μ*_11*s*_, *μ*_22*s*_) = corr(*μ*_12*s*_, *μ*_22*s*_) = *η*), the correlation matrix **C** is of the following structure,

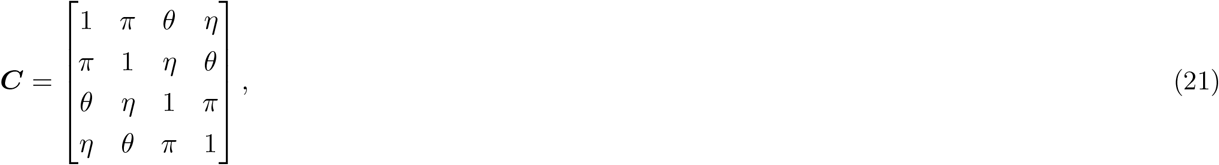

where the correlations *θ, η* and *π* are such that the correlation matrix **C** is positive semi-definite.

Now we derive the variance-covariance structure of the quad-variate (*τ*_1*s*_,*τ*_2*s*_,*λ*_1*s*_, *λ*_2*s*_)^*T*^ under the LME formulation (11). With the indicator variable *I_c_* defined in (10), the four variables are the varying intercepts and slopes and can be expressed as 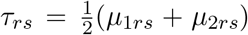 and *λ*_*rs*_ = *μ*_1*rs*_ – *μ*_2*rs*_ (*r* = 1,2). Furthermore, the correlation matrix for the quad-variate (*τ*_1*s*_, *τ*_2*s*_, *λ*_1*s*_, *λ*_2*s*_)^*T*^ is of the the following block diagonal form that justifies the independence assumption between the varying intercepts and varying slope in the LME formulation (11),

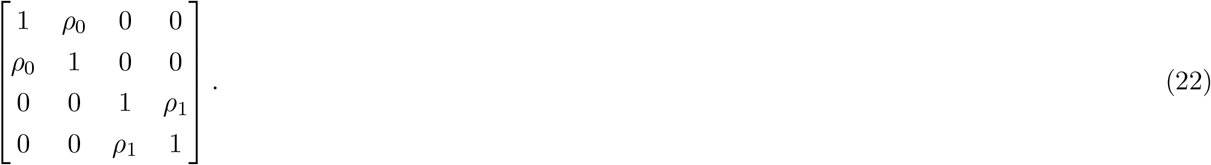

We obtain the correlation *ρ*_0_ between the two varying intercept components as

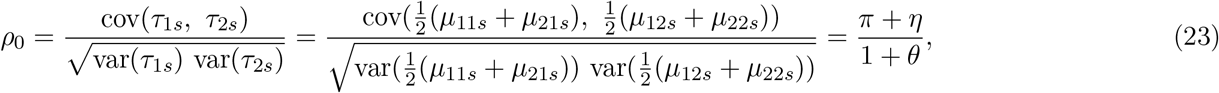

and the correlation *ρ*_1_ between the two varying slope components as

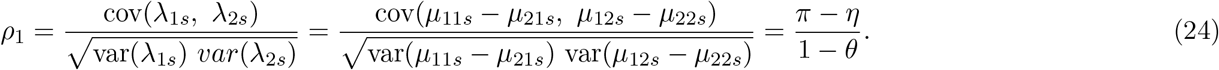

The correlation of 0s in (22) can be similarly derived as in (23) and (24).

## D Underestimation from the CLM-based method for a contrast between two conditions

The extent of underestimation by CLM follows a similar derivation for the case of a contrast between two conditions as that of a single condition. Based on the distribution assumption 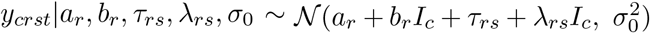 in the LME model (11) and the homoscedasticity assumption *σ*_λ_1__ = *σ*_λ_2__ = *σ*_λ_, we have

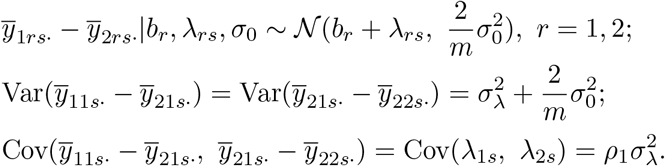

Plugging the above results into the definition of ICC for the contrast, we immediately see the degree to which the CLM generates lower estimates,

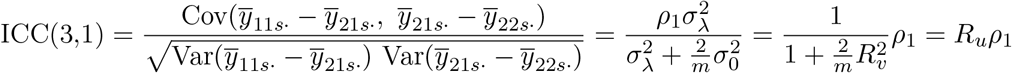

where 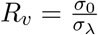 is the variability ratio and 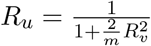 is the underestimation rate.

The CLM derived estimates can be corrected. If the cross-trial variability *σ*_0_ is known, we could restore the accuracy of ICC by removing the trial-related variance component 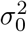 from the denominator of the ICC formulation,

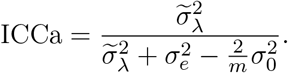

The underestimation for the average effect between the two conditions can be similarly derived. In fact, all the formulas remain the same as long as we replace the symbols *b_r_*, λ and *ρ*_1_ by *a_r_*, *τ* and *ρ*_0_.

## E Simulations of trial-level LME modeling for a contrast

Simulations for the reliability with a contrast under the LME model (11) are similar to the situation with a single condition but with a slightly more complexity. With the correlation matrix ***C*** in (21) across the four condition-level effects (*μ*_11*s*_, *μ*_12*s*_, *μ*_21*s*_, *μ*_22*s*_)^*T*^ and the homoscedasticity assumption in (20): *q*_11_ = *q*_21_ = *q*_12_ = *q*_22_ = 1, the cross-session reliability in the LME model (11) for the average and contrast between the two conditions can be simulated per the formulas (23) and (24). Below are the manipulation parameters:

1. 4 simulated reliability values *ρ*_1_ = 0.3, 0.5, 0.7 and 0.9 that, respectively, correspond to four sets of correlation structure **C** in (21): (*π, θ, η*) = (0.7, 0.5, 0.55), (0.7, 0.5, 0.45), (0.7, 0.5, 0.35), (0.7, 0.5, 0.25).
2. 4 different numbers of subjects: *n* = 20, 40, 100 and 200,
3. 4 different numbers of trials per session: *m* = 20, 40, 100 and 200,
4. 4 different ratios of cross-trial variability *σ*_0_ relative to cross-subject variability 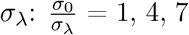 and 10,
5. 2 different sets of population-level effects across the two sessions, (*a*_11_, *a*_12_, *a*_21_, *a*_22_) = (0, 0, 0, 0) and (1.0, 0.9, 0, 0).
6. 3 different approaches to assessing cross-session reliability:

a. conventional ICC based ANOVA/LME (2) through aggregation across trials and conditions;
b. cross-session reliability *ρ*_1_ based on the LME formulation (11);
c. conventional ICC adjusted by removing the cross-trial variability 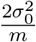 per formula (14).

Each of these 4 × 4 × 4 × 4 × 2 × 3 = 1536 combinations was simulated 1000 times. During each simulation, data were randomly drawn through the following steps:

1. For each subject *p*, obtain subject-level effects during the two sessions: 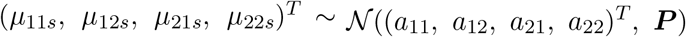, where ***P*** = ***C***.
2. Draw simulated data per the LME formulation (9): 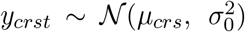, *c* = 1,2; *r* = 1,2; *s* = 1, 2,.., *n*; *t* = 1, 2,.., *m*.
3. Solve the two LME models, (2) and (11), using the function *lmer* in the *R* package `Ime4`.
4. Recover the simulated parameters including the three reliability estimates. Specifically, the conventional ICC is obtained through the formula (3) while the cross-session reliability *ρ*_1_ is estimated through the LME model (11). In addition, adjusted ICC is obtained per formula (14).

The simulation results largely follow a similar pattern to the situation for a single condition (Fig. 11).

## F Hyperpriors adopted for BML modeling

The prior distribution for all lower-level (e.g., subject) effects considered here is Gaussian, as specified in the respective model; for example, see the distribution assumptions in the BML models (15, 16, 17, 18). In addition, prior distributions (usually called hyperpriors) are needed for three types of model parameters in each model: (a) population effects or location parameters (e.g., population-level intercept and slopes), (b) standard deviations or scaling parameters for lower-level effects, and (c) various parameters such as the covariances in a variance-covariance matrix and the degrees of freedom in Student’s *t*-distribution. Noninformative hyperpriors are adopted for population-level effects. In contrast, weakly-informative priors are utilized for standard deviations of lower-level parameters such as varying intercepts and slopes at the subject level, and such hyperpriors include a Student’s half-t(3,0,1) or a half-Gaussian 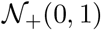 (a Gaussian distribution with restriction to the positive side of the respective distribution). For variance-covariance matrices, the LKJ correlation prior (Lewandowski et al., 2009) is used with the shape parameter taking the value of 1 (i.e., jointly uniform over all correlation matrices of the respective dimension). Lastly, the standard deviation *σ* for the residuals utilizes a half Cauchy prior with a scale parameter depending on the standard deviation of the input data. The hyperprior for the degrees of freedom, *ν*, of the Student’s *t*-distribution is Γ(2, 0.1). The consistency and full convergence of the Markov chains were confirmed through the split statistic 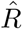 being less than 1.1 (Gelman et al., 2013). The effective sample size (or the number of independent draws) from the posterior distributions based on Markov chain Monte Carlo simulations was more than 200 so that the quantile (or compatibility) intervals of the posterior distributions could be estimated with reasonable accuracy.

**Figure 11:**
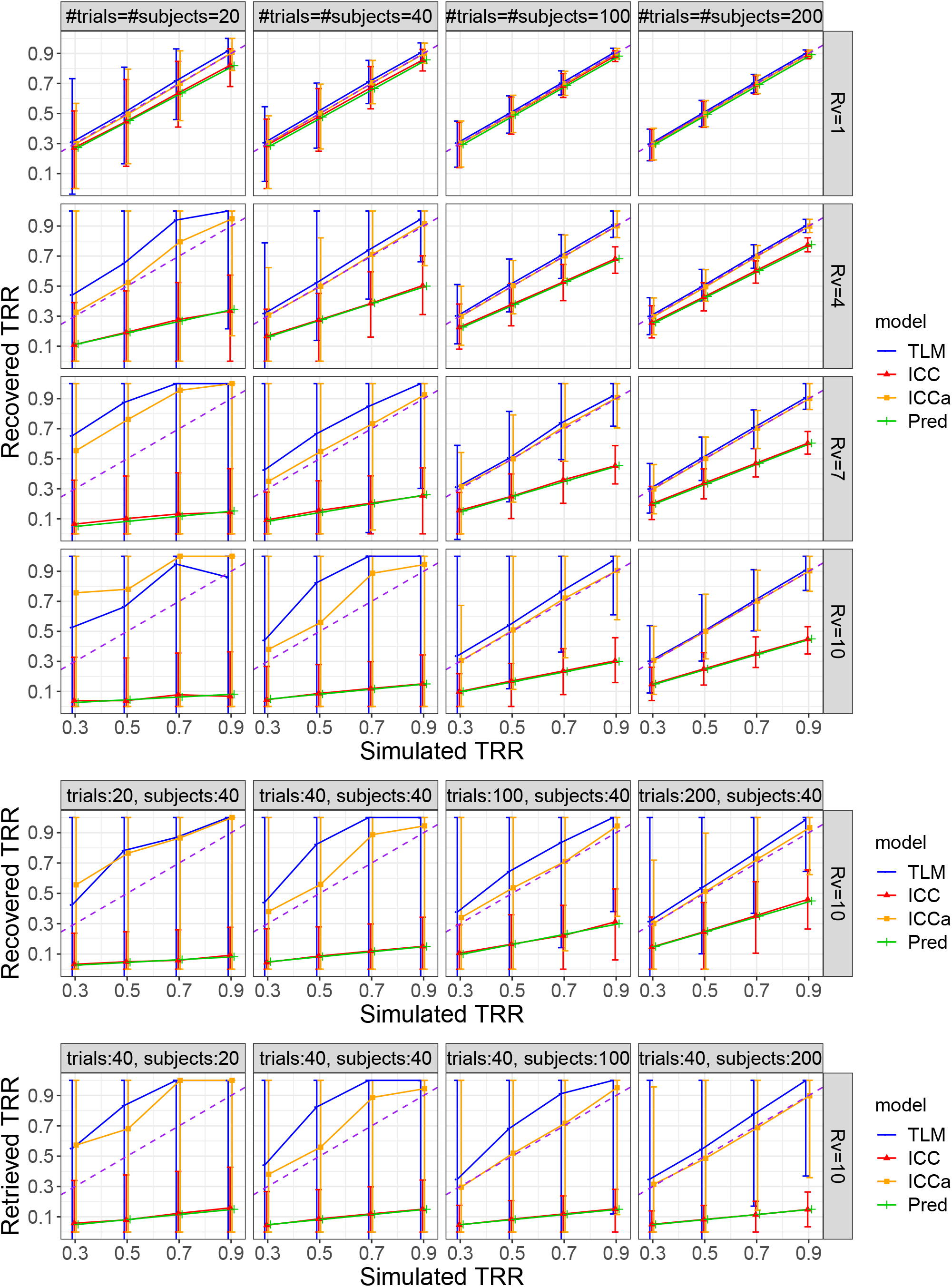
Simulation results for a contrast. The four columns correspond to the sample size of subjects and trials while the four rows are the varying standard deviation ratios of 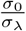. The *x*- and *y*-axis are the simulated and recovered reliability, respectively. Each data point is the median among the 1000 simulations with the error bar showing the 90% highest density interval. The dashed purple diagonal line indicates an idealized retrieval of the parameter value based on the data-generating model (11). The green line shows the theoretically predicted ICC per the formula (6).

## G Flanker Image acquisition and preprocessing

### Image Acquisition

Neuroimaging data were collected on a 3T GE Scanner using a 32-channel head coil across two separate sessions. After a sagittal localizer scan, an automated shim calibrated the magnetic field to decrease signal dropout from a susceptibility artifact. Echo-planar images were acquired across four functional runs at the following specifications: flip angle = 60°, echo time = 25 ms, repetition time = 2000 ms, 170 volumes per run, four runs, with an acquisition voxel size of 2.5 × 2.5 × 3 mm. The first 4 images from each run were discarded to ensure that longitudinal magnetization equilibrium was reached. Structural images were collected using a high-resolution T1-weighted magnetization-prepared rapid acquisition gradient echo (MPRAGE) sequence for co-registration with the functional data. Images were collected with a flip angle of 7° at a voxel size of 1 mm isotropic.

### Image Preprocessing

Neuroimaging data were analyzed using AFNI version 20.3.00 (http://afni.nimh.nih.gov/afni/; Cox, 1996) with standard preprocessing including despiking, slice-timing correction, distortion correction, alignment of all volumes to a base volume with minimum outliers, nonlinear registration to the MNI template, spatial smoothing with a 6.5mm FWHM kernel, masking, and intensity scaling. Final voxel size was 2.5 × 2.5 × 2.5 mm. We excluded any pair of successive TRs in which the sum head displacement (Euclidean norm of the derivative of the translation and rotation parameters) between those TRs exceeded 1 mm. TRs in which more than 10% of voxels were outliers were also excluded. Participants’ datasets were excluded if the average motion per TR after censoring was greater than 0.25 mm or if more than 15% of TRs were censored for motion or outliers. In addition, 6 head motion parameters were included as nuisance regressors in individual-level models.

## Footnotes

1 Instead of using the indicator variable *I_c_* in (10) to represent the two conditions, one may adopt the following dummy coding (as in Haines et al., 2020): one condition is coded as 1 while the other as 0 (reference condition). As a result, the slopes in the LME model (11) still correspond to the contrast; however, the intercepts are associated with the reference condition.

2 https://rb.gy/uakhmj

3 The reliability analysis scripts used in this paper are available at https://github.com/afni-gangc/TRR

4 https://surfer.nmr.mgh.harvard.edu/optseq/

5 The aggregation step more accurately reflects the typical preprocessing in behavior data than in neuroimaging. The following two pipelines are not strictly commutative in fMRI data analysis: (a) obtain condition-level effect estimates from time series regression with one regressor per condition, and (b) perform time series regression with one regressor per trial and then obtain the condition-level effect estimate by averaging the trial-level regression coefficients. The aggregation step in our simulations follows the latter pipeline as a rough approximation for the former.

